# Longitudinal network analysis links multi-omics data with clinical outcomes in Parkinson’s disease

**DOI:** 10.64898/2026.11.17.767355

**Authors:** Marina Vargas-Fernández, Pedro Carmona-Sáez, Jordi Martorell-Marugán

**Affiliations:** GENYO. Centre for Genomics and Oncological Research: Pfizer / University of Granada / Andalusian Regional Government, PTS Granada, Granada 18016, Spain; Department of Statistics and Operational Research, University of Granada, Granada, 18071, Spain; Computational Neuroscience. Joint Unit in Biomedical Imaging and Artificial Intelligence FISABIO–CIPF. Foundation for the Promotion of Health and Biomedical Research in the Valencian Region (Fisabio), 46024 Valencia, Spain

**Author notes:** Correspondence to: Jordi Martorell-Marugán, PhD, Foundation for the Promotion of Health and Biomedical Research in the Valencian Region (Fisabio). Computational Neuroscience. Joint Unit in Biomedical Imaging and Artificial Intelligence FISABIO– CIPF. Carrer d’Eduardo Primo Yúfera 3, Valencia 46024, Spain. Correspondence may also be addressed to: Professor Pedro Carmona-Sáez, PhD, University of Granada, Department of Statistics and Operational Research. Avenida de la Fuente Nueva S/N, Granada 18071, Spain. These authors contributed equally to this work (Pedro Carmona-Sáez, Jordi Martorell-Marugán).

## Abstract

Parkinson’s disease (PD) shows heterogeneous progression of motor and non-motor symptoms. Although transcriptomic and proteomic alterations have been reported in blood, plasma and cerebrospinal fluid (CSF), their associations with longitudinal clinical trajectories remain unclear. We integrated longitudinal multi-omics and clinical data from 2,606 participants harmonized by the Accelerating Medicines Partnership for Parkinson’s Disease (AMP-PD). We used MariNET, a novel network analysis framework for heterogeneous data based on linear mixed-effects models, to examine associations between molecular features and disease progression.

In blood, we found modest associations of oxidative phosphorylation, proteasome-mediated protein catabolism and T-helper 1 cytokine production with motor, non-motor and cognitive scores at the transcriptomic level. In CSF, dopa decarboxylase (DDC) levels were associated with treatment-related motor complications, measured by Part IV of the Unified Parkinson’s Disease Rating Scale, and prolactin (PRL) levels with daytime sleepiness. DDC and PRL associations had the same direction in plasma.

Our findings link biological processes and proteins to specific clinical domains of PD across repeated observations. Our framework is applicable to other longitudinal multi-omics studies and freely available as a module in the MariNET R package.

## Introduction

Parkinson’s disease (PD) is a progressive neurodegenerative disorder with motor and non-motor symptoms. Motor symptoms, including bradykinesia, rigidity, tremor and postural instability, arise primarily from dopaminergic neuron loss in the substantia nigra. Cognitive impairment, sleep disturbances, autonomic dysfunction, mood disorders and sensory deficits are also common, indicating biological processes beyond dopaminergic neurodegeneration [1].

Symptomatic treatments such as levodopa and dopamine agonists improve motor function but do not prevent neurodegeneration or halt disease progression. Clinical presentation and progression rates vary substantially among patients, suggesting that multiple molecular mechanisms contribute to disease evolution [2, 3].

High-throughput molecular profiling has enabled systematic characterization of PD-associated molecular alterations [4, 5]. Large-scale transcriptomic and proteomic studies of peripheral fluids have revealed perturbations in pathways involving mitochondrial dysfunction, neuroinflammation, protein aggregation, vesicle trafficking and immune responses [6]. Cerebrospinal fluid (CSF) provides a proximal view of central nervous system pathology, whereas peripheral blood offers a minimally invasive window into systemic alterations, including inflammatory and immune responses [7, 8].

However, variation in circulating immune cell composition also shapes blood transcriptomic signals and can confound disease-associated differential expression. The Parkinson’s Progression Markers Initiative (PPMI), a large international, multicenter observational study of early-stage, untreated PD, reported enrichment of neutrophil degranulation and related immune pathways in PD blood samples. These findings underscore the need to account for cell type composition in whole-blood transcriptomic analyses [9].

Despite these advances, links between molecular alterations and the progression of specific clinical manifestations remain unclear. Most molecular studies compare patients with controls or assess progression using global outcomes. It therefore remains uncertain which molecular changes are associated with longitudinal trajectories in motor impairment, cognition, sleep, olfaction and treatment-related complications.

Repeated clinical assessments offer an opportunity to address this gap. However, integrating them with high-dimensional molecular data requires accounting for technical and biological variability while preserving the longitudinal structure. Existing network-based approaches, such as Weighted Gene Co-expression Network Analysis (WGCNA), are typically applied to cross-sectional molecular data and do not directly model longitudinal associations between molecular features and clinical trajectories.

Here, we integrated longitudinal clinical assessments with transcriptomic and proteomic data from the Accelerating Medicines Partnership for Parkinson’s Disease (AMP-PD) cohort [10]. We used MariNET, a network framework based on linear mixed-effects models, to examine molecular–clinical associations while accounting for repeated measurements and relevant covariates. By combining whole-blood transcriptomics with plasma and CSF proteomics, we aimed to identify biological processes and molecular markers associated with specific clinical domains in PD.

## Results

### Study overview and data description

The AMP-PD initiative integrates longitudinal clinical and multi-omics data from several PD cohorts. We analyzed whole-blood transcriptomics and CSF and plasma proteomics linked to repeated clinical assessments (Figure 1).

**Figure 1.**
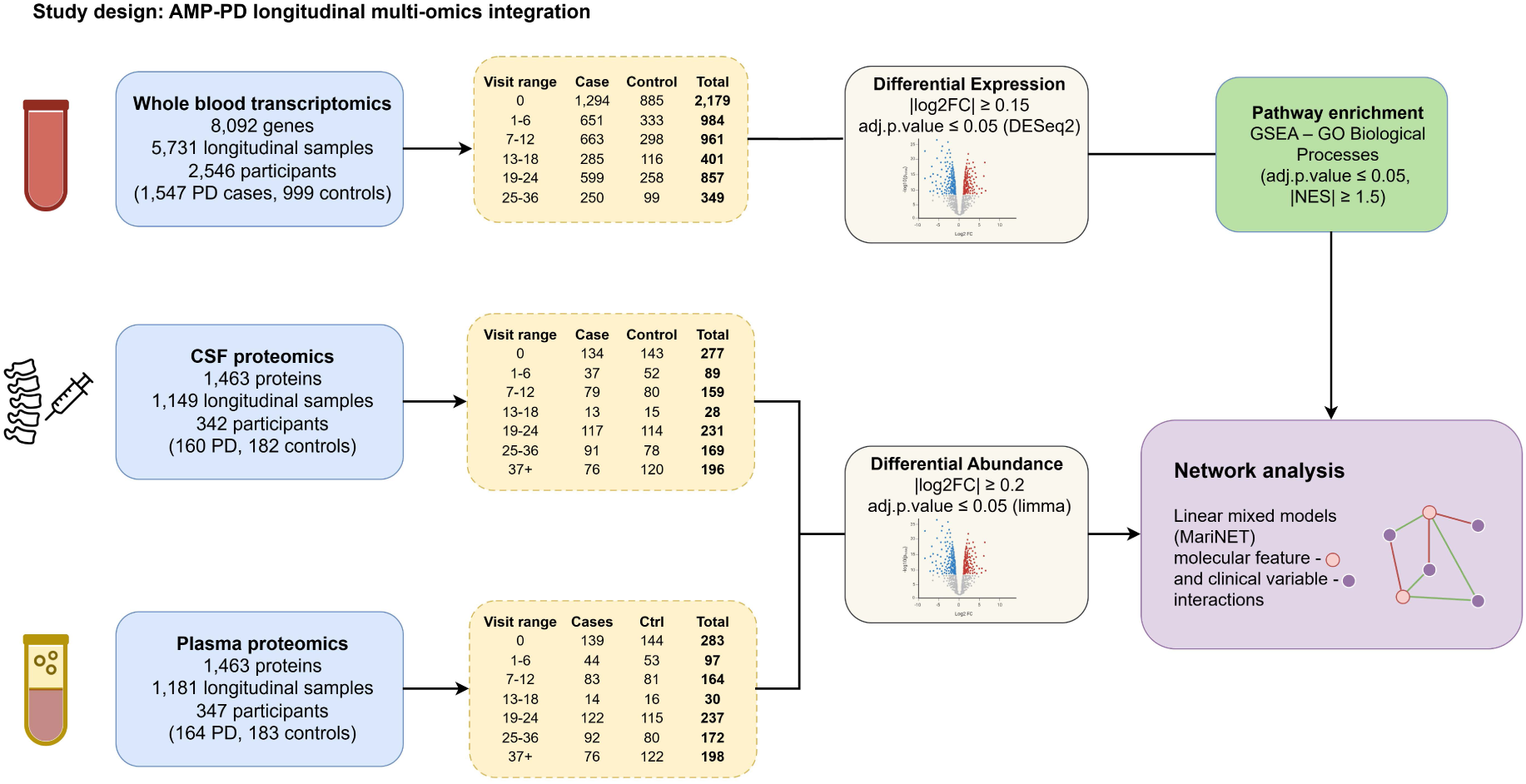
Overview of the study design. Longitudinal transcriptomics, CSF proteomics and plasma proteomics datasets from AMP-PD were processed independently for differential expression and abundance analyses. Prioritized molecular features were then integrated with longitudinal clinical data using network analysis based on linear mixed-effects models.

After quality control and sample harmonization, the transcriptomics dataset comprised 5,731 RNA-seq samples from 2,546 participants (1,547 PD cases and 999 controls), with 8,092 genes retained for analysis. Participants had a mean age of 62.2 ± 11.0 years; 54.9% were male and 93.5% were White.

The CSF proteomics dataset comprised 1,149 longitudinal samples from 342 <u>participants</u> (160 PD cases and 182 controls), and the plasma dataset comprised 1,181 samples from 347 participants (164 PD cases and 183 controls). In each biofluid, 1,463 proteins were quantified using four Olink panels. Both cohorts had a mean age of 62.6 ± 10 years and were 64% male.

### Differential expression and abundance analyses prioritize features for network analyses

To prioritize molecular features for network analysis, we performed case–control differential expression analyses for transcriptomics and differential abundance analyses for proteomics independently in each dataset. We considered the longitudinal design by including visit month as covariate in the linear models.

Cell-type composition is a major source of variation in whole-blood transcriptomics and can confound disease-associated signals if not properly controlled [11, 12]. To prevent this effect from driving differential expression and downstream analyses, we incorporated cell-type composition as a covariate in the generalized linear models. However, direct hematology measurements were available for only 1,371 of the 6,547 samples retained at this stage.

Following the approach of [9], which addressed the same limitation in AMP-PD transcriptomics data, we imputed neutrophil percentages for the remaining 5,176 samples (79.1%) using a linear regression model trained on a 31-gene neutrophil marker signature (adjusted R² = 0.809). We included the resulting estimates as covariates in the differential expression models to account for neutrophil-driven variation and better isolate PD-associated signals.

In addition to cell-type composition, we evaluated the impact of biological and technical variables on whole-blood transcriptomics by fitting linear models between each covariate and the principal components of the expression data. Visit month, sex, age, race, neutrophil percentage, sequencing plate, RNA integrity number (RIN), and normalization volume were associated with the leading principal components (Supplementary Figure 1, Supplementary Table 1). Among highly correlated technical variables (concentration, submitted volume, and normalization volume; |ρ| ≥ 0.99), normalization volume explained the largest proportion of expression variance and was therefore retained. RNA input and specimen quantity were excluded because their effects were captured by the remaining covariates (Supplementary Table 3). After adjustment, these associations were removed while the association with diagnosis was preserved (Supplementary Figure 2, Supplementary Table 2), and the selected variables were included in the differential expression models.

Using this approach, we identified 13 differentially expressed genes (DEGs) in whole-blood transcriptomics data (Figure 2A, Supplementary Table 4). FOLR3, encoding a neutrophil-enriched folate-binding protein, was the only significantly upregulated gene, whereas the remaining 12 DEGs were downregulated, including H1-0, BPGM, XK, and CA1 among the most strongly altered.

**Figure 2.**
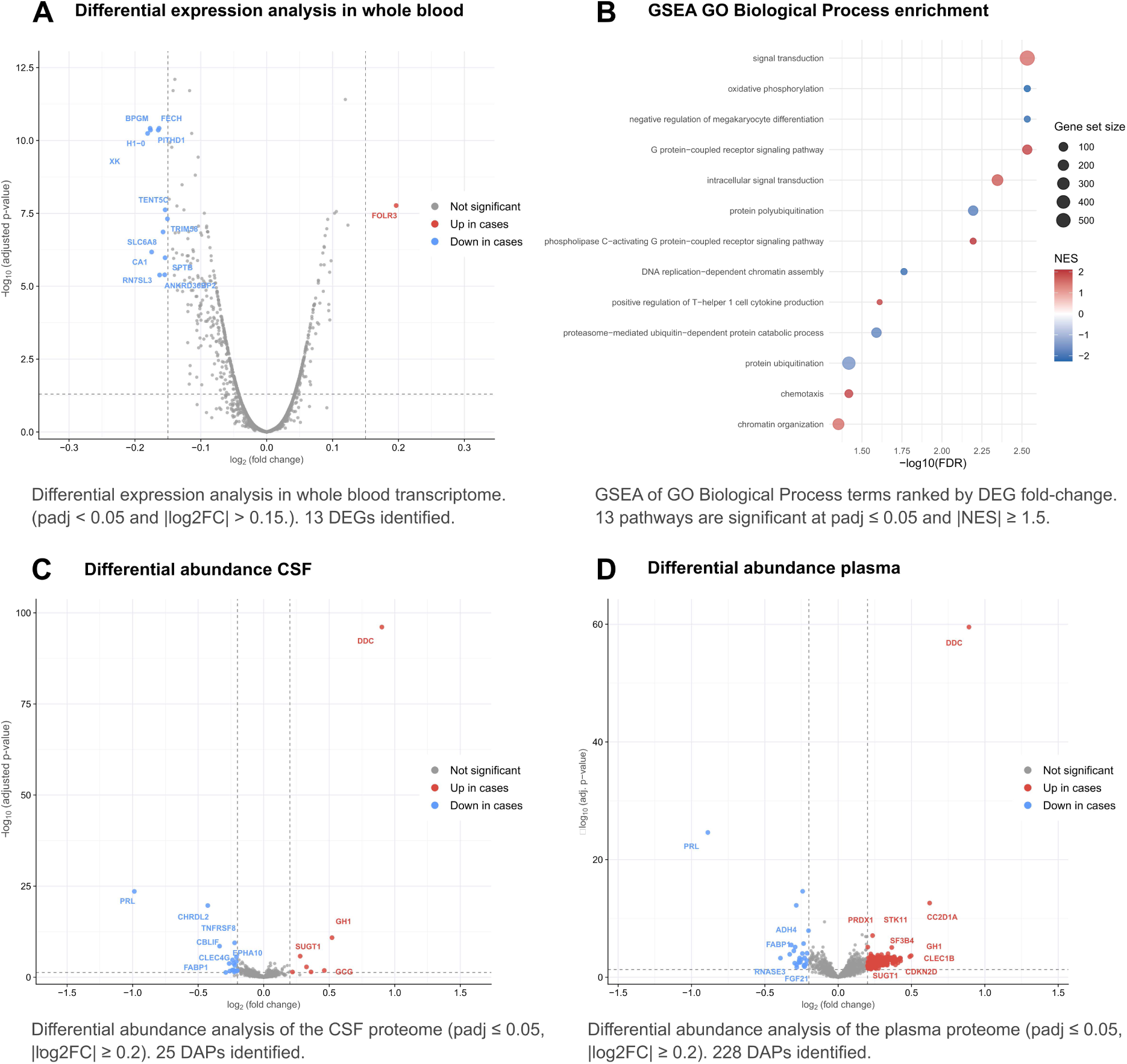
Differential expression and abundance analyses across transcriptomics and proteomics datasets in PD. **(A)** Volcano plot of case-control DEGs in whole blood. **(B)** Significantly enriched GO BP terms identified by GSEA. **(C)** Volcano plot of case-control DAPs in CSF. **(D)** Volcano plot of case-control DAPs in plasma.

To facilitate biological interpretation beyond individual genes and generate network-level features, we performed Gene Set Enrichment Analysis (GSEA) using Gene Ontology Biological Process (GO BP) terms. We identified 16 significantly enriched terms (FDR < 0.05, |NES| ≥ 1.5), which were reduced to 13 after removing redundancy (Figure 2B, Supplementary Table 5). Negatively enriched processes in PD included oxidative phosphorylation (NES = −2.26), negative regulation of megakaryocyte differentiation (NES = −2.17), DNA replication-dependent chromatin assembly (NES = −2.13), and protein polyubiquitination (NES = −1.86). Positively enriched processes included phospholipase C-activating G protein-coupled receptor signaling (NES = 2.10), G protein-coupled receptor signaling (NES = 1.95), and chemotaxis (NES = 1.95).

For the proteomics datasets, no technical covariates were available. Therefore, we evaluated the effects of biological variables using the same approach applied to transcriptomics (Supplementary Figures 3-6, Supplementary Tables 6-9). Based on these analyses, we included sex and age as covariates in the differential abundance models.

We identified 25 differentially abundant proteins (DAPs) in CSF (Figure 2C, Supplementary Table 10) and 228 in plasma (Figure 2D, Supplementary Table 11). In CSF, prolactin (PRL), a pituitary hormone whose secretion is inhibited by dopamine, and dopa decarboxylase (DDC), the enzyme that converts levodopa into dopamine, showed the largest abundance changes. Other strongly altered proteins included somatotropin (GH1), pro-glucagon (GCG), the precursor of glucagon and GLP-1, and lactoperoxidase (LPO), an antimicrobial peroxidase.

In plasma, most DAPs were increased in PD (202 upregulated and 26 downregulated proteins). DDC and PRL again showed the largest abundance changes, followed by SMAD1, PDLIM7, and SRC, a tyrosine kinase belonging to a family known to phosphorylate α-synuclein. Five proteins were differentially abundant in both CSF and plasma and changed in the same direction: DDC, GH1, and suppressor of G2 allele of SKP1 homolog (SUGT1) were increased in PD, whereas PRL and fatty acid-binding protein 1 (FABP1), an intracellular lipid transport protein, were decreased.

### Blood transcriptomics reveals pathway-level associations with clinical outcomes

We first examined associations between PD-associated DEGs and clinical outcomes (Figure 3A, Supplementary Table 12). Significant associations included a negative relationship between TENT5C expression and olfactory performance (UPSIT score). FOLR3, the only upregulated DEG, was negatively associated with non-motor experiences of daily living (UPDRS1) and positively associated with PDQ-39 mobility (Supplementary Table 12).

**Figure 3.**
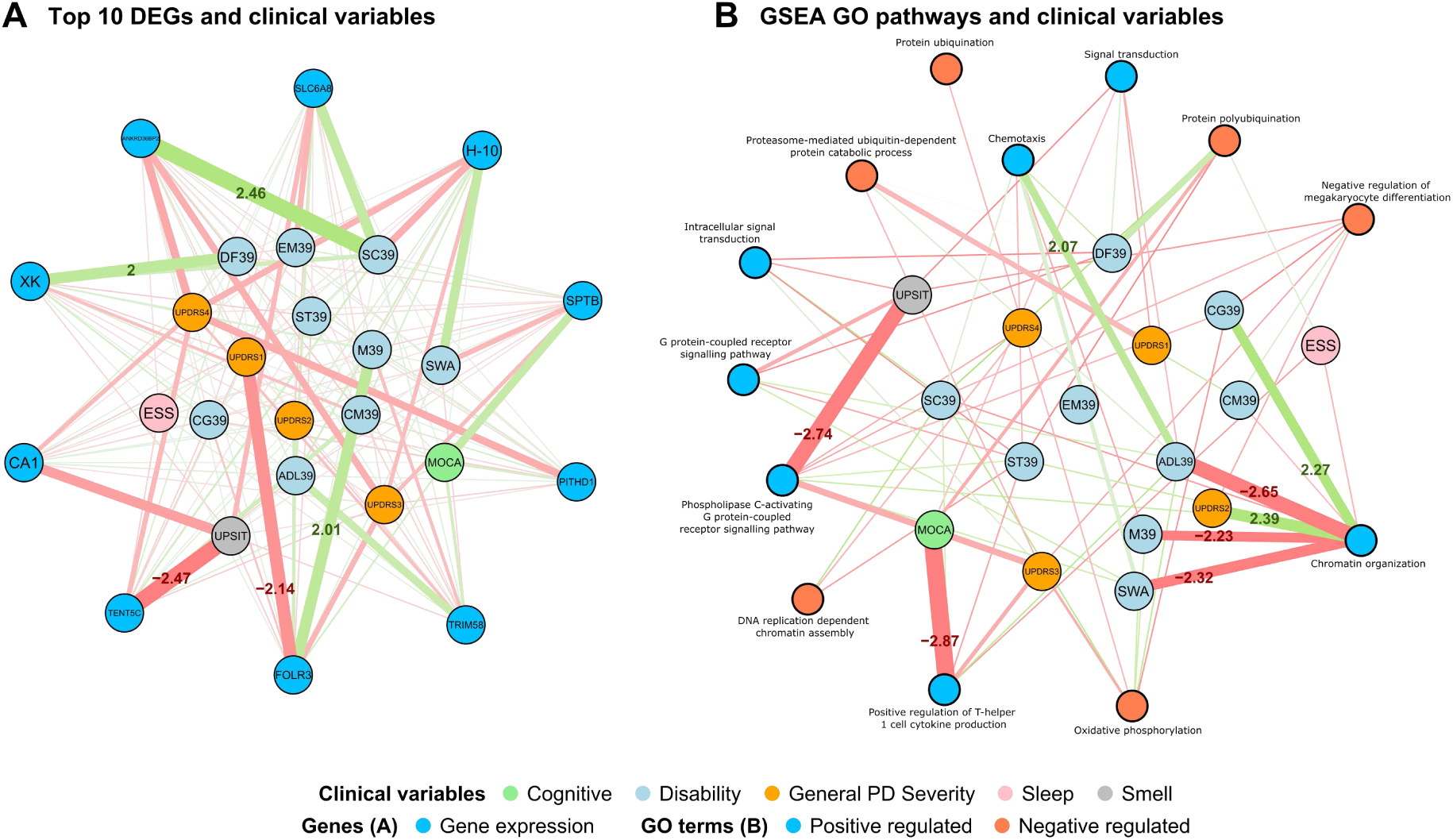
Longitudinal MariNET networks linking whole-blood transcriptomic features to clinical variables in PD. **(A)** Gene–clinical network for the 10 differentially expressed genes with the highest maximum |t|. **(B)** Pathway-clinical network for the significantly enriched GO BP terms, using sample-level pathway activity scores estimated with GSVA. Blue and orange pathway nodes indicate terms upregulated and downregulated in PD, respectively. Green edges indicate positive associations, red edges negative associations, and edge width is proportional to |t|. Edge labels show t-values for associations with |t| > 2. Clinical nodes are coloured by domain and placed in the same positions in both panels. Abbreviations: ESS, Epworth Sleepiness Scale; UPDRS1-4, Movement Disorder Society Unified Parkinson’s Disease Rating Scale Parts I–IV; MoCA, Montreal Cognitive Assessment; PDQ-39, Parkinson’s Disease Questionnaire-39 (domains: M39, Mobility; ADL39, Activities of Daily Living; EM39, Emotional Well-being; ST39, Stigma; SC39, Social Support; CG39, Cognition; CM39, Communication; DF39, Bodily Discomfort); SWA, Modified Schwab and England Activities of Daily Living Scale; UPSIT, University of Pennsylvania Smell Identification Test.

We next examined pathway-level associations using GSVA activity scores derived from GO BP terms significantly enriched by GSEA (Figure 3B, Supplementary Table 13). Selected PD-relevant pathways are summarized in Table 1. Among immune-related pathways, positive regulation of T-helper 1 cell cytokine production was negatively associated with cognitive performance, whereas chemotaxis was positively associated with PDQ-39 activities of daily living. Within the ubiquitin-proteasome system, protein polyubiquitination was positively associated with PDQ-39 bodily discomfort, while proteasome-mediated ubiquitin-dependent protein catabolic process was negatively associated with UPDRS1. We also observed weaker associations between oxidative phosphorylation and UPDRS3, and between G protein-coupled receptor signaling and UPSIT.

**Table 1.** Top associations between clinically relevant GO BP terms and clinical variables. For each GO BP term, we report the clinical variable with the largest absolute t-value in the MariNET network. Positive NES values indicate enrichment among genes upregulated in PD.

| GO BP term | Positive/Negative regulation (NES) | Interaction sign (t-value) | Top clinical variable |
|---|---|---|---|
| Oxidative phosphorylation | -2.26 | -1.63 | UPDRS3 |
| Proteasome-mediated ubiquitin-dependent protein catabolic process | -1.76 | -1.85 | UPDRS1 |
| Protein polyubiquitination | -1.86 | 1.97 | Discomfort39 |
| G protein-coupled receptor signaling pathway | 1.95 | -1.75 | UPSIT |
| Positive regulation of T-helper 1 cell cytokine production | 1.95 | -2.87 | MOCA |
| Chemotaxis | 1.95 | 2.07 | ADL39 |

### CSF proteomics identifies clinically relevant molecular associations

Longitudinal network analysis linking CSF proteomics to clinical variables identified associations with both motor and non-motor features of PD (Figure 4, Supplementary Table 14). DDC abundance was positively associated with UPDRS4 (t = 2.80), whereas PRL was negatively associated with the ESS score (t = −2.85). CBLIF was positively associated with UPDRS1 and negatively associated with Cognition39 and UPDRS2. CGC was positively associated with Mobility39.

**Figure 4.**
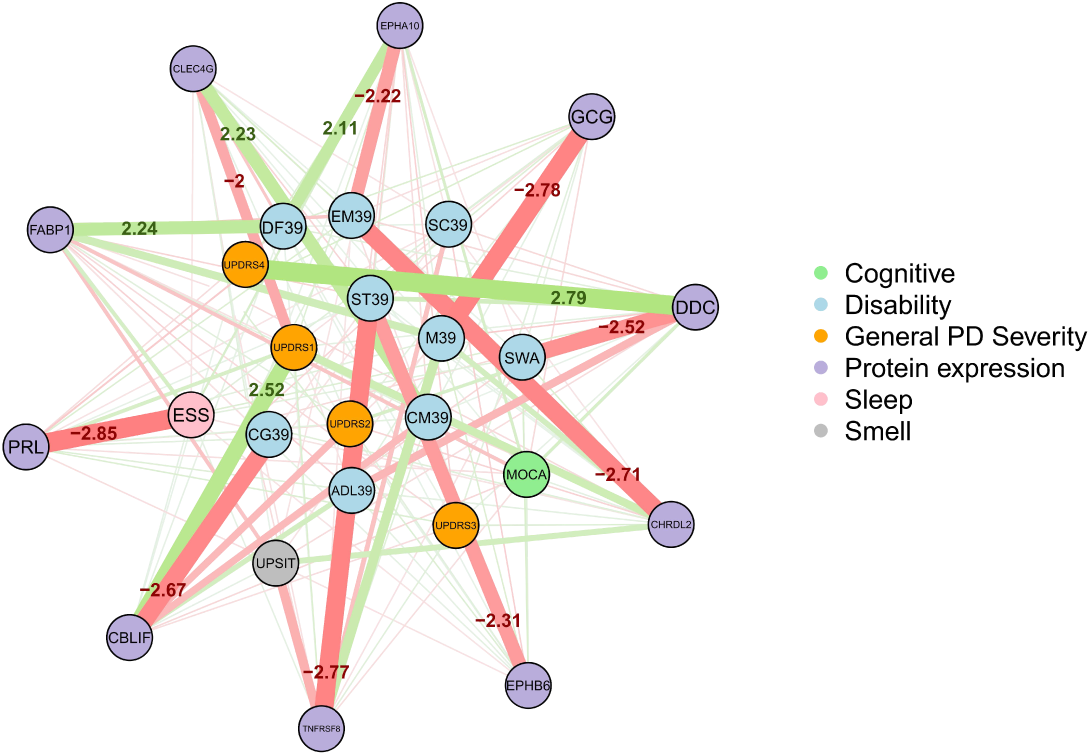
Interaction network between CSF protein abundance and PD symptomatology. Green and red edges indicate positive and negative associations, respectively, and edge width is proportional to |t|. Edge labels are shown for associations with |t| > 2.

To further characterize the DDC-UPDRS4 association, we examined UPDRS4 scores in the complete-case CSF cohort (415 visits from 149 participants; PD cases only). UPDRS4 was markedly zero-inflated: 174 visits (41.9%) scored 0, whereas only 8 (1.9%) scored ≥13. Treatment-related complications (UPDRS4 > 0) were present in 241 visits (58.1%).

Higher CSF DDC abundance was associated with increased odds of treatment-related complications (OR = 3.55, 95% CI [2.00, 6.32], p = 1.6 × 10⁻⁵ after adjustment for treatment status; unadjusted OR = 3.70, p = 2.9 × 10⁻⁵). The predicted probability of complications increased from <10% at low DDC levels to nearly 100% at high levels (Figure 5A). Because 98.1% of visits occurred under treatment, adjustment for treatment status had little effect on the estimated association. No DDC × visit month interaction was detected (β ≈ 0, 95% CI [−0.044, 0.044], p = 0.999), indicating a stable association over time.

**Figure 5.**
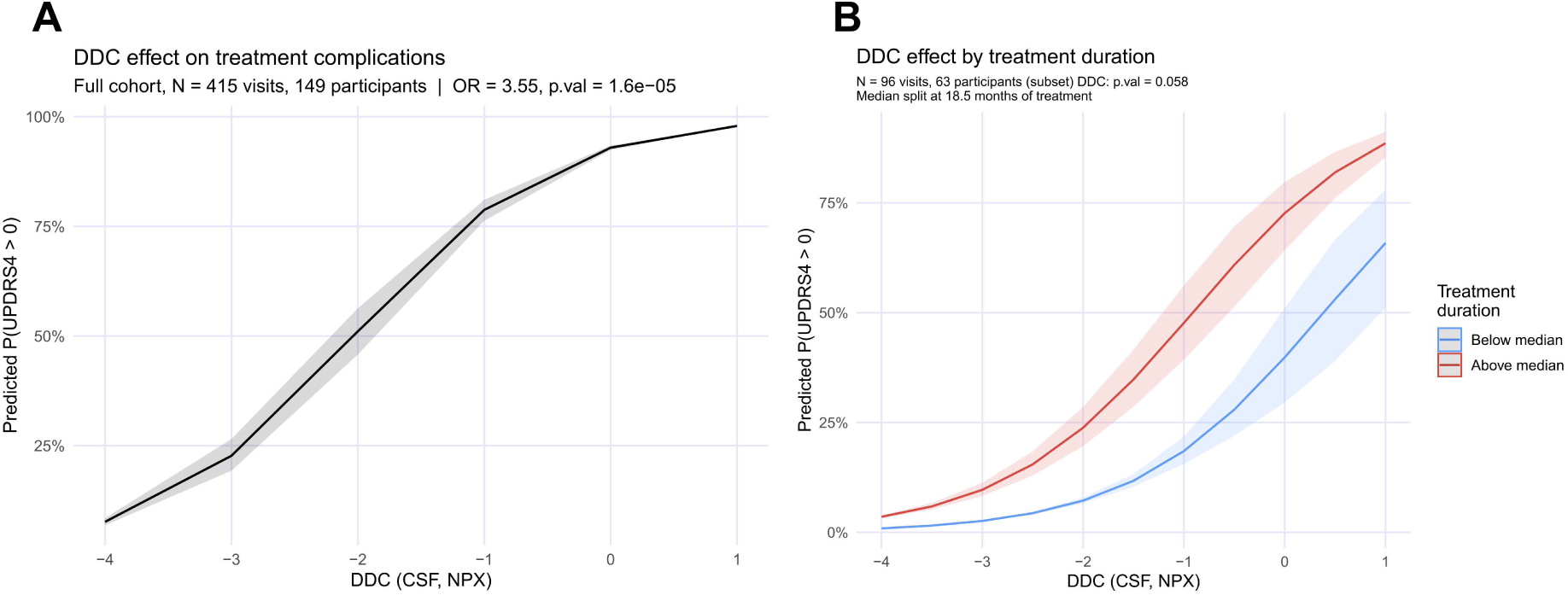
CSF DDC levels predict treatment-related complications in PD. DDC levels are reported as Olink Normalized Protein eXpression (NPX) units. **(A)** Model-estimated probability of motor complications (UPDRS4 > 0) across CSF DDC levels in the full cohort. Shaded areas indicate 95% confidence intervals. **(B)** The same relationship stratified by treatment duration (median split, 18.5 months) in participants with documented treatment start dates.

In visits with a documented treatment start date (96 visits from 63 participants), the association remained similar after adjustment for treatment duration (OR = 2.91, p = 0.058), although statistical support was inconclusive. Treatment duration was independently associated with complications (OR ≈ 1.07 per month, p = 0.035). Predicted probabilities were consistently higher above than below the median treatment duration (18.5 months), while preserving the overall DDC-complication relationship (Figure 5B).

Plasma proteomics partially reflects proteomics-clinical associations in CSF Differential abundance analysis identified 228 plasma DAPs between PD cases and controls. We then constructed a longitudinal network linking plasma proteomics to clinical variables and visualized the 10 most connected proteins (Figure 6A, Supplementary Table 15). Major associations included STK11 with UPDRS3, DDC with UPDRS4, consistent with the CSF network, PRL with ESS, SF3B4 with UPSIT, PRDX1 with Cognition39 and UPSIT, and CDKN2D with Mobility39 and Discomfort39.

**Figure 6.**
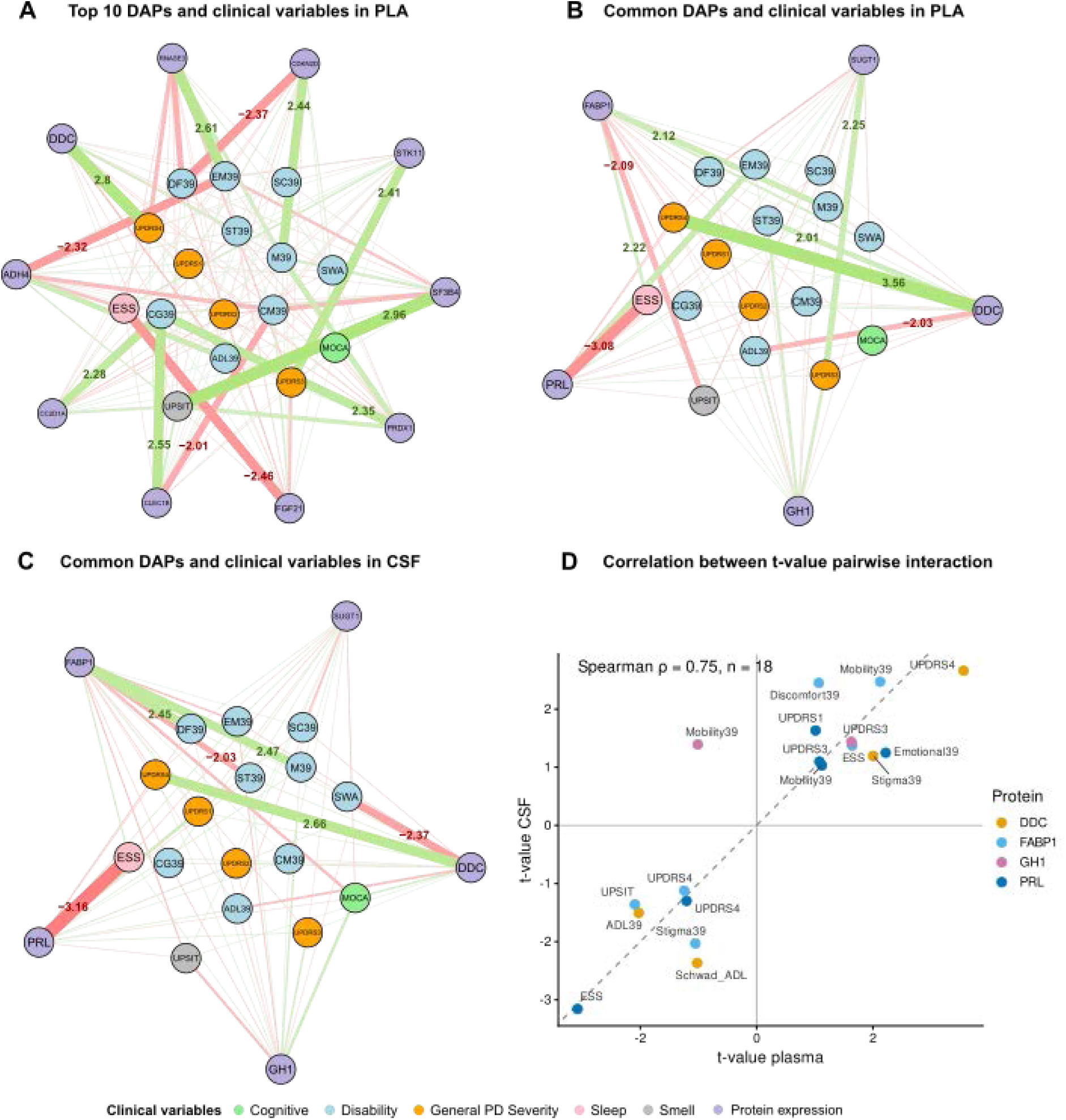
Longitudinal MariNET networks linking plasma and CSF proteomics to clinical variables in PD. **(A)** Plasma protein-clinical network showing the 10 DAPs with the highest maximum |*t*|. **(B)** Plasma network for the five proteins differentially abundant in both plasma and CSF (DDC, PRL, FABP1, GH1, and SUGT1). **(C)** Corresponding CSF network. Green and red edges indicate positive and negative associations, respectively, and edge width is proportional to |*t*|. Edge labels are shown for associations with |*t*| > 2. **(D)** Correlation between plasma and CSF *t*-values for the five shared proteins. Points are coloured by protein, the dashed line indicates identical values in both biofluids, and the Spearman correlation is shown (ρ = 0.75, *n* = 18 pairs with |*t*| > 1 in both biofluids; SUGT1 reached no associations above this threshold).

To assess cross-biofluid consistency, we generated proteomics-clinical networks for the five proteins differentially abundant in both CSF and plasma (DDC, PRL, GH1, FABP1, and SUGT1) using measurements from each biofluid (Figure 6B-C; Supplementary Table 16). DDC was positively associated with UPDRS4 in both CSF and plasma (t = 2.661 and 3.560, respectively) and negatively associated with Schwab & England ADL and ADL39, with stronger effects in CSF. PRL was negatively associated with ESS in both biofluids (t = −3.158 and −3.078). FABP1 showed positive associations with Mobility39 and Discomfort39, and negative associations with UPSIT and Stigma39, in both CSF and plasma.

We next compared t-values for all protein-clinical pairs between the plasma and CSF networks. Among 80 evaluated pairs, 18 reached |t| > 1 in both biofluids, of which 17 showed concordant directions of association (Spearman ρ = 0.75, p < 0.001, Figure 6D). Similar agreement was observed when all pairs were considered (Spearman ρ = 0.63, p < 0.001). The only discordant association was GH1-Mobility39, which was positive in CSF and negative in plasma. GH1 remained positively associated with UPDRS3 in both biofluids, whereas no SUGT1 association reached the |t| > 1 threshold.

## Discussion

Identifying relationships between longitudinal omics and clinical data remains challenging. Here, we adapted our network methodology, MariNET, to integrate longitudinal molecular profiles with repeated clinical assessments in PD. By linking longitudinal molecular profiles with repeated clinical assessments in PD, our method highlighted genes, pathways and proteins associated with specific domains in PD. Unlike case-control designs, our approach models each molecular feature against multiple clinical scales simultaneously, linking molecular signals to specific clinical domains rather than to disease status alone. Linear mixed models allow all visits per participant to be included while accounting for within-participant correlation, increasing power relative to single-timepoint analyses. The framework also allows adjustment for technical and biological covariates.

The associations found at transcriptional level in whole blood revealed several genes with potential links to PD related processes, although their interpretation should be approached cautiously, as circulating transcripts may reflect systemic responses rather than central nervous system mechanisms. Whole blood RNA-seq signal is additionally shaped by variation in circulating immune cell composition, which can itself be altered in PD and can confound disease-associated differential expression if left uncorrected [9]. To address this, we predicted neutrophil composition and incorporated it as a covariate in the differential expression models used here. Even so, FOLR3, the only upregulated gene, is highly expressed in neutrophils, so its identified association with UPDRS1 may partly reflect residual compositional signal.

Among the significant DEGs, TENT5C expression showed a strong association with UPSIT scores. TENT5C (FAM46C) is a non-canonical poly(A) polymerase involved in B-cell biology and immunoglobulin regulation [13]. This association may reflect either an immune-related peripheral signal or an unrecognized neuronal role. In addition, H1-0, a linker histone, was among the most strongly downregulated genes, and chromatin assembly was also downregulated at the pathway level. Given the involvement of chromatin remodeling and epigenetic dysregulation in neurodegeneration [14, 15], altered H1-0 expression may reflect broader chromatin changes associated with PD. The network analysis reported a positive association between H1-0 expression and activities of daily living (Schwad_ADL), further linking H1-0 expression to preserved functional status.

The pathways activity at transcriptional level was more informative and consistent with mechanisms previously implicated in PD. Oxidative phosphorylation was downregulated in PD and its activity was inversely associated with motor symptom severity (UPDRS3), in line with the mitochondrial dysfunction described in PD [16, 17]. The ubiquitin-proteasome pathways, which regulate protein degradation, were also downregulated, and proteasome-mediated protein catabolism was inversely associated with UPDRS1. Impaired protein degradation contributes to α-synuclein accumulation in PD [18, 19]. The observed association is compatible with a relationship between altered proteasome-related activity and non-motor symptom burden, although the present analysis cannot establish a causal contribution. Among the upregulated pathways, T-helper 1 cytokine production was associated with lower cognitive performance and chemotaxis with worse daily-living scores, in line with reports linking peripheral immune markers to clinical progression in PD [20].

At proteomics level, the most relevant result was the association of DDC, the most differentially abundant protein in PD, with treatment-related complications measured by the UPDRS-IV score. DDC catalyzes the conversion of levodopa to dopamine and is therefore central to PD treatment. Peripheral DDC activity can reduce levodopa availability in the brain, which is why DDC inhibitors such as carbidopa are co-administered [21]. In this context, higher DDC levels could reduce the effective availability of levodopa in the central nervous system. DDC level in CSF has also been recently proposed as a diagnostic biomarker in Lewy body disorders [22].

Further analyses supported the DDC association after adjustment for treatment status and found no evidence that it changed across follow-up. In the smaller subset with treatment-duration data, the estimated association remained in the same direction and of comparable magnitude, although the evidence was inconclusive (p = 0.058). This may reflect variability in levodopa metabolism contributing to the pulsatile dopaminergic stimulation implicated in motor fluctuations and dyskinesias [23]. Notably, we also observed this association in plasma. However, treatment is a prerequisite for motor complications and nearly all visits were on treatment, so treatment may itself increase DDC rather than DDC driving complications. Furthermore, binary treatment status has little variance in this cohort, treatment duration was available for only a subset of visits and underestimates exposure in participants treated before enrolment, and no levodopa dose was available.

PRL secretion is negatively regulated by dopamine [24–26]. Despite dopaminergic neuron loss in PD, PRL levels were lower in cases than controls, consistent with the suppressive effects of dopaminergic medication. Likewise, the negative association between PRL and daytime sleepiness may partly reflect treatment effects [27]. Together with DDC, PRL may therefore capture treatment exposure in addition to disease biology.

CBLIF encodes gastric intrinsic factor, the glycoprotein required for intestinal absorption of vitamin B12 (cobalamin). Lower CSF vitamin B12 status has been associated with faster motor progression in early, untreated PD, particularly greater decline in gait and mobility [28]. Vitamin B12 has also been shown to directly interact with α-synuclein, inhibiting its fibrillogenesis and reducing amyloid-induced cytotoxicity in vitro [29]. In our analysis, CBLIF was positively associated with UPDRS1 and negatively associated with cognitive aspects of quality of life (Cognition39) and UPDRS2, raising the possibility that altered intrinsic factor levels (and vitamin B12 availability) could contribute to non-motor and cognitive symptom burden through modulation of α-synuclein aggregation, although this link remains speculative and would require direct B12 measurement to confirm.

GCG encodes preproglucagon, the precursor of glucagon and glucagon-like peptide-1 (GLP-1). GLP-1 receptor agonists such as exenatide have been investigated as disease-modifying treatments in PD, with a recent phase 3 trial reporting effects on motor progression [30]. In our analysis, GCG was negatively associated with mobility (Mobility39), consistent with a role for the glucagon/GLP-1 signaling axis in motor function, although causality cannot be inferred from the present data.

Regarding plasma proteomics, several of the top interacting proteins point toward mitochondrial and oxidative-stress-related mechanisms, complementing the oxidative phosphorylation pathway association identified in the transcriptomic analysis. STK11 (LKB1), an upstream regulator of AMPK signaling, was positively associated with motor severity and is linked to mitochondrial homeostasis and neuroprotection in PD [31]. STK11 was positively associated with motor severity (UPDRS3) in our plasma network. PRDX1, an antioxidant protein implicated in neurodegenerative disease [32], was positively associated with cognitive performance and olfactory function.

To examine whether clinical associations observed in CSF were also present in plasma, we focused on the proteins differentially abundant in both biofluids (DDC, PRL, FABP1, GH1 and SUGT1). Among the 18 protein–clinical pairs with |t| > 1 in both networks, 17 showed the same direction of association, and their t-values were strongly correlated between fluids. This agreement extended across motor, functional, quality-of-life, sleep and olfactory domains. DDC, PRL and FABP1 showed fully concordant associations, whereas GH1 showed a single divergent and weak association. SUGT1 showed cognitive and social associations in CSF that were not reproduced in plasma; more broadly, no cognitive association was conserved across fluids. These differences suggest that some cognitive correlates may be more specific to the brain environment. Overall, the concordance supports further investigation of plasma as a minimally invasive source of these molecular correlates.

Taken together, CSF and plasma provide both overlapping and complementary molecular information: shared DAPs largely share their clinical associations, while the DAPs specific to each biofluid point to central processes (e.g., neuroinflammation, dopaminergic dysfunction and treatment-related mechanisms) in CSF, and to peripheral correlates of motor and functional progression in plasma. The identification of proteins with consistent associations across both compartments highlights their potential as biomarkers accessible through minimally invasive sampling. Furthermore, although repeated measurements were modelled using mixed-effects models, the network analysis was not designed to separate within-participant changes from between-participant differences. The identified associations therefore reflect longitudinal relationships across repeated observations rather than evidence that molecular changes track or predict individual progression. Replication in longitudinal cohorts with dose information and pre-treatment sampling would be needed before these proteins can be proposed as biomarkers.

## Methods

### Blood transcriptomics data processing and analysis

A total of 8,461 whole-blood RNA-seq samples were available from the AMP-PD initiative (BioFIND, PDBP and PPMI cohorts). Samples with more than 5% unmapped or mismatched reads were excluded (7,244 retained), followed by samples with an RNA integrity number (RIN) below 6.5, leaving 6,547 samples and 58,780 genes in the raw count matrix. Analyses were restricted to participants with a single, consistent diagnosis of PD or healthy control, excluding those diagnosed with other neurological disorders (5,790 samples). Genes expressed at >10 counts per million (CPM) in at least 80% of samples were retained (8,191 genes), and 99 mitochondrial, ribosomal and globin genes were removed, leaving 8,092 genes. Finally, samples whose combination of sequencing plate and race was represented by a single sample were excluded (n = 59) to avoid inestimable cells in the design matrix. The final dataset comprised 5,731 samples (3,742 from PD cases and 1,989 from controls) from 2,546 participants.

In order to account for the confounding effect of blood cell composition on gene expression, we adapted the approach previously developed by Xu et al. [9], who demonstrated that incorporating the neutrophil percentage as covariate effectively adjusts this bias. Hematology measurements were available for 1371 (20.9%) samples all from the PPMI cohort. In these samples, a linear regression model was fitted to predict measured neutrophil percentage from the variance-stabilized (VST) expression of a previously defined 31-gene neutrophil marker signature [9], (adjusted R² = 0.80924). This model was then used to impute neutrophil percentage for the remaining 5,176 samples (79.1%).

We used the swamp R package to assess the influence of technical and biological covariates on the first ten principal components of the TMM-normalized gene expression. (Supplementary Figure 1, Supplementary Table 1). The contribution of each variable was summarized as the proportion of expression variance it explained, computed as its R² with each component weighted by the variance explained by that component. Because nearly all variables reached significance at this sample size, covariates were selected based on variance explained rather than p-values. To avoid overcorrection, only normalization volume was selected among a set of strongly correlated technical variables (Spearman |ρ| ≥ 0.99), which was the one explaining the most variance was retained (Supplementary Table 2). Sequencing plate was fully nested within cohort, which was not modelled separately (Supplementary Figure 2, Supplementary Table 2). After adjustment, variance explained by the remaining technical variables was ≤ 0.25%, except for well position (2.1%). Well position was not included, as its per-component R² was of the order expected by chance for a 94-level factor, and modelling it would have required 93 additional coefficients per gene. The association with diagnosis was maintained after correction (Supplementary Table 3).

Differential expression analysis was performed with DESeq2 [33] on raw counts, using normalization factors calculated with the trimmed mean of M values (TMM) method [34]. The design formula was:

~~~
design ∼ Condition + Visit month + Sex + Race + Age + Neutrophil (%) + Sequencing plate + RIN + Normalization volume
~~~

Controls were set as the reference level, so positive log₂ fold changes indicate higher expression in PD cases. Continuous covariates were standardized before model fitting. Dispersions were estimated with the standard DESeq2 workflow using a local regression fit. Differentially expressed genes (DEGs) were defined by a Benjamini-Hochberg adjusted p-value < 0.05 and |log₂ fold change| > 0.15.

To investigate pathway-level perturbations in transcriptomic data, we performed gene set enrichment analysis (GSEA) focusing on Gene Ontology (GO) biological process term [35] using the fgsea package [36]. Terms with at least five genes in the ranked list were tested with the multilevel algorithm, and terms with an adjusted p-value < 0.05 and an absolute normalized enrichment score (NES) ≥ 1.5 were considered significantly enriched. Redundant GO terms were removed with the collapsePathways function of fgsea.

Sample-level pathway activity scores for the non-redundant terms were computed with the GSVA method through the getScores function of pathMED [37]. Scores were calculated on the log-CPM expression matrix after removing the effect of the differential expression covariates with removeBatchEffect. These scores were used as pathway nodes in the MariNET network analysis.

### CSF and plasma proteomics data processing and analysis

Proteomics data for both cerebrospinal fluid (CSF) and plasma were obtained from the Accelerating Medicines Partnership for Parkinson’s Disease (AMP-PD) initiative. These data were generated using four targeted Olink 1536 proteomics panels (i.e. cardiometabolic, inflammation, neurology and oncology), which we merged for our analyses. Quality control (QC) and preprocessing was performed by the AMP-PD consortium. This included the removal of samples and analytes failing detection thresholds, transformation of abundance values, normalization and statistical correction for technical batch effects.

Differential abundance analysis between PD patients and controls was performed independently in CSF and plasma. The analysis was conducted using the limma package in R [38]. A linear model was fitted to the data for each protein with the following design:

~~~
Design ∼ 0 + Condition + Visit month + Sex + Age
~~~

The intercept was removed to allow direct estimation of group-level means for each condition, enabling explicit contrast specification (Case vs. Control) via makeConstrasts() [39].

Following the model fit (lmFit), the contrast of interest (PD vs. Control) was calculated. Empirical Bayesian moderation was then applied (eBayes) to stabilize the variance estimates. Proteins meeting |log₂(FC)| ≥ 0.2 and adjusted p-value ≤ 0.05 were considered differentially abundant proteins (DAPs) for network analyses. UniProt accessions were mapped to HGNC gene symbols via Ensembl BioMart.

### Processing of clinical data

The clinical data associated with the omics datasets includes demographic variables such as sex, race, diagnosis, medical history, medication, environmental risk factors, and disease staging based on the Hoehn and Yahr scale. For the network analyses, we used continuous summary scores from key clinical assessments, including the Unified Parkinson’s Disease Rating Scale (UPDRS) Parts I, II, III, and IV; the Parkinson’s Disease Questionnaire (PDQ-39); the Schwab and England Activities of Daily Living (ADL) Scale; the Montreal Cognitive Assessment (MoCA); the Epworth Sleepiness Scale (ESS); and the University of Pennsylvania Smell Identification Test (UPSIT).

Temporal entries were excluded if they lacked observations for at least 70% of the selected variables. Identical measurements were omitted. Repeated measurements where all variables had identical values at different time points (e.g., baseline and month 0 due to study definitions) were collapsed into a single observation.

### Multimodal integration and network analysis

Associations between omics data and clinical variables were estimated with MariNET [40], a network framework for longitudinal data. In MariNET, each variable is modeled as the outcome of a linear mixed-effects model (LMM) with the remaining variables as fixed effects and participant identifier and grouping variables as random intercept, which accounts for repeated measurements and intragroup correlation. Each variable is a node, and the edge between two nodes is weighted by the t-value of the corresponding fixed-effect coefficient. The sign of the edge therefore indicates the direction of the association, whereas its absolute weight reflects the magnitude of the corresponding *t*-statistic, which depends on both the estimated coefficient and its standard error.

Applied directly to molecular data, the original approach would model every feature against every other, allowing gene–gene associations to dominate the network and obscure the relationships of interest. We therefore modified the model structure to focus on associations between clinical variables and molecular features (Figure 7). Each molecular feature was modeled as a function of the clinical variables alone, without other molecular features as predictors. Each clinical variable was modeled as a function of the molecular features and the remaining clinical variables, so that its association with each molecular feature was adjusted for the other clinical variables. Edge weights were interpreted as measures of model-based statistical association, rather than as direct measures of effect size or biological importance.

**Figure 7:**
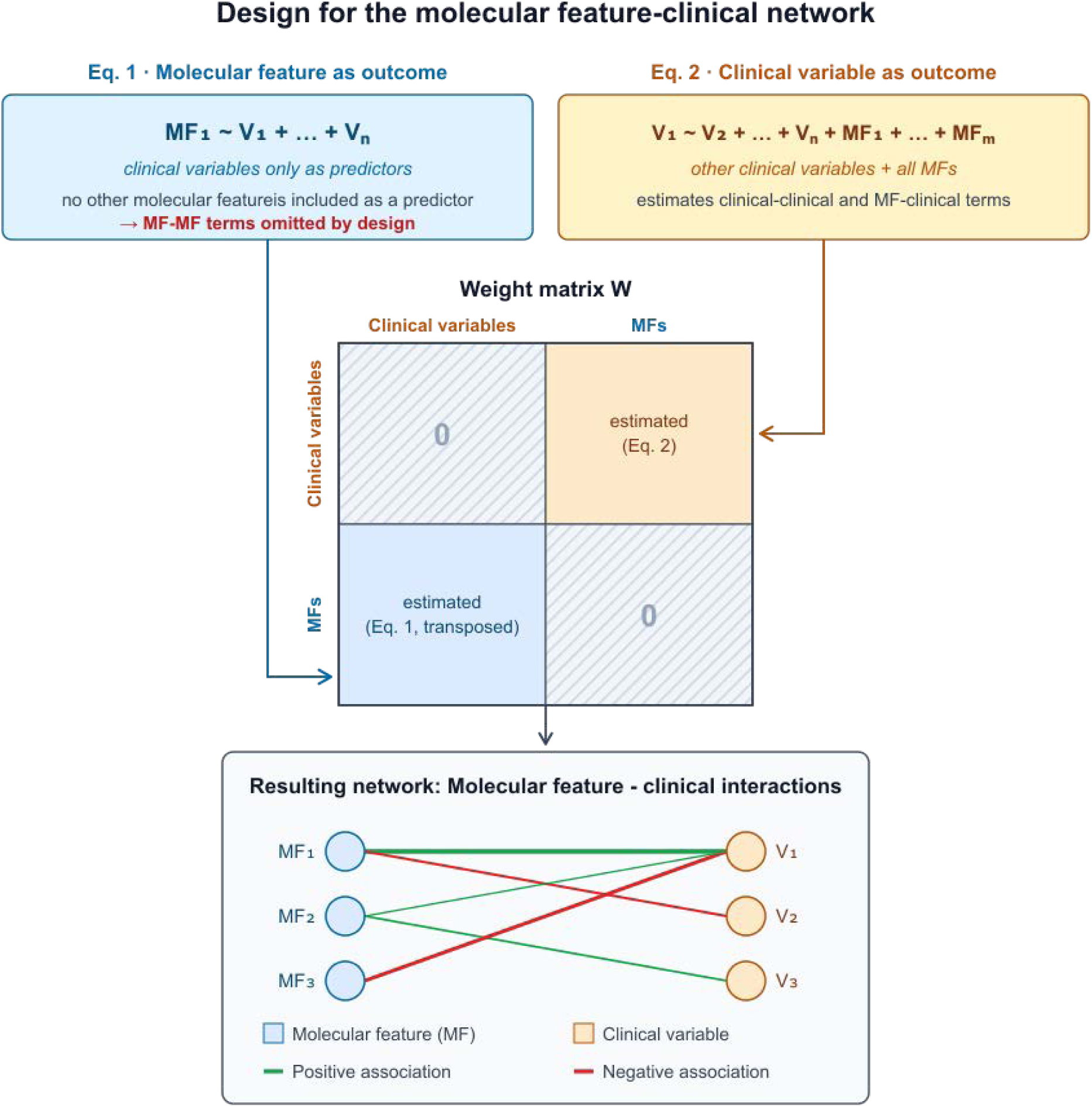
Network construction strategy. Each molecular feature (MF) is modeled as a function of clinical variables (Eq. 1), and each clinical variable as a function of the MFs and the remaining clinical variables (Eq. 2). MF-MF associations are not modeled, and clinical-clinical coefficients, although estimated in Eq. 2, are excluded from the network, which therefore contains only MF-clinical edges.

For visualization, network plots display the top 10 molecular features ranked by their maximum absolute t-value across clinical variables.

Each omics layer was analyzed in a separate network with the same set of clinical variables. In all cases, molecular data were matched to clinical assessments by participant identifier and time point, preserving the longitudinal design.

For transcriptomics, the DEGs identified in whole-blood RNA-seq were included as molecular nodes using normalized expression. For proteomics, the DAPs identified in CSF and plasma were included in separate networks for each fluid type using normalized abundance. For pathway-level analysis, sample-wise activity scores were computed with GSVA and were included as molecular feature nodes.

Network analyses were restricted to PD cases, to focus on associations between molecular features and clinical outcomes within patients rather than on case-control differences. Patients with idiopathic and non-idiopathic PD were analyzed as a single group. The final dataset comprised 3,414 visits from 1,443 participants.

### DDC and treatment complications association

The association between CSF DDC and UPDRS Part IV (UPDRS4) identified in the network was further studied. The analysis was restricted to PD cases with complete data for DDC, UPDRS4, age at baseline and sex (415 visits from 149 participants). DDC abundance was used as Olink NPX values, without standardization. The distribution of UPDRS4 scores in this sample was first examined descriptively. Because scores were zero-inflated, which violates the Gaussian assumption of the linear mixed models used in network construction, the outcome was dichotomized as presence (UPDRS4 > 0) or absence of motor complications.

A logistic mixed-effects model was fitted with DDC abundance, age at baseline, sex and treatment status as fixed effects, and participant identifier as a random intercept to account for repeated visits. Treatment status was defined per visit as “yes” if the participant was on levodopa, a dopamine agonist or any other PD medication. The model was also fitted without treatment status to obtain the unadjusted estimate. To assess whether the association changed over follow-up, a DDC × visit month interaction (visit month in months from baseline) was added, with a random slope for visit month per participant, falling back to a random intercept only if the fit was singular. This model was compared with the model without the interaction by likelihood-ratio test.

To evaluate cumulative dopaminergic exposure as an alternative explanation, a sensitivity analysis was performed in the subset of visits with a documented treatment start date. Treatment duration (months since treatment initiation, truncated at 0) replaced treatment status as a continuous covariate, with age, sex and the participant random intercept retained. For visualization, predicted probabilities of motor complications across DDC abundance were obtained from this model separately for visits above and below the median treatment duration. Effects are reported as odds ratios with Wald 95% confidence intervals. Models were fitted with glmer (lme4, binomial family, logit link).

## Data availability

The data analyzed in this research was obtained from the Accelerating Medicine Partnership® (AMP®) Parkinson’s Disease (AMP PD) Knowledge Platform. Data used for this paper can be accessed by demand. The datasets used correspond to v4.0 of ‘Transcriptomics Data’, ‘Targeted Proteomics Data’, and ‘Clinical Assessment Data’.

## Code availability

The code implemented for this analysis can be accessed on https://github.com/marinavargasf/Omics_Integration.

## Acknowledgments

Data used in the preparation of this article were obtained from the Accelerating Medicines Partnership® (AMP®) Parkinson’s Disease (AMP PD) and Parkinson’s Disease & Related Disorders (AMP PDRD) Knowledge Platform Knowledge Platform. For up-to-date information on the study, visit https://www.amp-pdrd.org.

The AMP® PD program is a public-private partnership managed by the Foundation for the National Institutes of Health and funded by the National Institute of Neurological Disorders and Stroke (NINDS) in partnership with the Food and Drug Administration (FDA), National Institute on Aging (NIA), Aligning Science Across Parkinson’s (ASAP) initiative; Celgene Corporation, a subsidiary of Bristol-Myers Squibb Company; GlaxoSmithKline plc (GSK); The Michael J. Fox Foundation for Parkinson’s Research (MJFF); AbbVie Inc.; Pfizer Inc.; Sanofi US Services Inc.; and Verily Life Sciences LLC.

The AMP® PDRD program is a public-private partnership managed by the Foundation for the National Institutes of Health and funded by the National Institute of Neurological Disorders and Stroke (NINDS) in partnership with the Food and Drug Administration (FDA), National Institute on Aging (NIA), AbbVie Inc.; Aligning Science Across Parkinson’s (ASAP) initiative; C2N Diagnostics, LLC; CurePSP; GlaxoSmithKline plc (GSK); Denali Therapeutics Inc.; Laboratory Corporation of America Holdings (Labcorp); The Michael J. Fox Foundation for Parkinson’s Research (MJFF); Sanofi US Services Inc.; and Verily Life Sciences LLC.

ACCELERATING MEDICINES PARTNERSHIP and AMP are registered service marks of the U.S. Department of Health and Human Services.

Clinical data and biosamples used in preparation of this article were obtained from the (i) Michael J. Fox Foundation for Parkinson’s Research (MJFF) and National Institute of Neurological Disorders and Stroke (NINDS) BioFIND study, (ii) NINDS Parkinson’s Disease Biomarkers Program (PDBP), and (iii) MJFF Parkinson’s Progression Markers Initiative (PPMI). Data were obtained from AMP PD release 4.0, downloaded on 18 November 2023.

BioFIND is sponsored by The Michael J. Fox Foundation for Parkinson’s Research (MJFF) with support from the National Institute for Neurological Disorders and Stroke (NINDS). The BioFIND Investigators have not participated in reviewing the data analysis or content of the manuscript. For up-to-date information on the study, visit www.michaeljfox.org/biofind.

The Parkinson’s Disease Biomarker Program (PDBP) consortium is supported by the National Institute of Neurological Disorders and Stroke (NINDS) at the National Institutes of Health. A full list of PDBP investigators can be found at https://pdbp.ninds.nih.gov/policy. The PDBP investigators have not participated in reviewing the data analysis or content of the manuscript.

PPMI is sponsored by The Michael J. Fox Foundation for Parkinson’s Research and supported by a consortium of scientific partners, found at https://www.ppmi-info.org/about-ppmi/who-we-are/study-sponsors. The PPMI investigators have not participated in reviewing the data analysis or content of the manuscript. For up-to-date information on the study, visit www.ppmi-info.org.

This work was supported by grant PID2024-156297OB-I00 funded by MICIU/AEI/10.13039/501100011033/ FEDER, UE. JMM was supported by Grant RYC2024-049532-I funded by MICIU/AEI/10.13039/501100011033 and by ‘ESF+’.

## Author contributions

MVF: Conceptualization, Formal analysis, Investigation, Methodology, Software, Visualization, Writing - original draft, writing - review & editing; PCS: Conceptualization, Supervision, Funding acquisition, Resources, Supervision, Writing - review & editing; JMM: Conceptualization, Data Curation, Funding Acquisition, Supervision, Validation, Writing - review & editing.

## References

[1] Radad K, Moldzio R, Krewenka C, Kranner B, Rausch W-D. Pathophysiology of non-motor signs in Parkinson’s disease: some recent updating with brief presentation. Exploration of Neuroprotective Therapy. 2023;3(1):24–46. doi:10.37349/ent.2023.00036

[2] Lee AJ, Kim C, Park S, Joo J, Choi B, Yang D, Jun K, Eom J, Lee S-J, Chung SJ, et al. Characterization of altered molecular mechanisms in Parkinson’s disease through cell type–resolved multiomics analyses. Science Advances. 9(15):eabo2467. doi:10.1126/sciadv.abo2467

[3] Poewe W, Seppi K, Tanner CM, Halliday GM, Brundin P, Volkmann J, Schrag A-E, Lang AE. Parkinson disease. Nature Reviews Disease Primers. 2017;3(1):17013. doi:10.1038/nrdp.2017.13

[4] Chang D, Nalls MA, Hallgrímsdóttir IB, Hunkapiller J, van der Brug M, Cai F, Kerchner GA, Ayalon G, Bingol B, Sheng M, et al. A meta-analysis of genome-wide association studies identifies 17 new Parkinson’s disease risk loci. Nature Genetics. 2017;49(10):1511–1516. doi:10.1038/ng.3955

[5] Morris HR, Spillantini MG, Sue CM, Williams-Gray CH. The pathogenesis of Parkinson’s disease. The Lancet. 2024;403(10423):293–304. doi:10.1016/S0140-6736(23)01478-2

[6] Imam F, Saloner R, Vogel JW, Krish V, Abdel-Azim G, Ali M, An L, Anastasi F, Bennett D, Pichet Binette A, et al. The Global Neurodegeneration Proteomics Consortium: biomarker and drug target discovery for common neurodegenerative diseases and aging. Nature Medicine. 2025;31(8):2556–2566. doi:10.1038/s41591-025-03834-0

[7] Sinha A, Srivastava N, Singh S, Singh AK, Bhushan S, Shukla R, Singh MP. Identification of differentially displayed proteins in cerebrospinal fluid of Parkinson’s disease patients: A proteomic approach. Clinica Chimica Acta. 2009;400(1):14–20. doi:10.1016/j.cca.2008.09.026

[8] Esteves AR, Cardoso SM. Differential protein expression in diverse brain areas of Parkinson’s and Alzheimer’s disease patients. Scientific Reports. 2020;10(1):13149. doi:10.1038/s41598-020-70174-z

[9] Xu KY, Violich I, Hutchins E, Alsop E, Nalls MA, Moore A, Blauwendraat C, Raphael Gibbs J, Cookson MR, Van Keuren-Jensen K, et al. Decreased SNCA expression in whole-blood RNA analysis of Parkinson’s disease adjusting for neutrophils. npj Parkinson’s Disease. 2025;11(1):292. doi:10.1038/s41531-025-01062-4

[10] Iwaki H, Leonard HL, Makarious MB, Bookman M, Landin B, Vismer D, Casey B, Gibbs JR, Hernandez DG, Blauwendraat C, et al. Accelerating Medicines Partnership: Parkinson’s Disease. Genetic Resource. Movement Disorders. 2021;36(8):1795–1804. doi:10.1002/mds.28549

[11] Xie X, Liu M, Zhang Y, Wang B, Zhu C, Wang C, Li Q, Huo Y, Guo J, Xu C, et al. Single-cell transcriptomic landscape of human blood cells. National Science Review. 2020;8(3):nwaa180. doi:10.1093/nsr/nwaa180

[12] Muñoz-Delgado L, Macías-García D, Periñán MT, Jesús S, Adarmes-Gómez AD, Bonilla Toribio M, Buiza Rueda D, Jiménez-Jaraba M del V, Benítez Zamora B, Díaz Belloso R, et al. Peripheral inflammatory immune response differs among sporadic and familial Parkinson’s disease. npj Parkinson’s Disease. 2023;9(1):12. doi:10.1038/s41531-023-00457-5

[13] Bilska A, Kusio-Kobiałka M, Krawczyk PS, Gewartowska O, Tarkowski B, Kobyłecki K, Nowis D, Golab J, Gruchota J, Borsuk E, et al. Immunoglobulin expression and the humoral immune response is regulated by the non-canonical poly(A) polymerase TENT5C. Nature Communications. 2020;11(1):2032. doi:10.1038/s41467-020-15835-3

[14] Labbé C, Lorenzo-Betancor O, Ross OA. Epigenetic regulation in Parkinson’s disease. Acta Neuropathologica. 2016;132(4):515–530. doi:10.1007/s00401-016-1590-9

[15] Zhang D, Zhang J, Wang Y, Wang G, Tang P, Liu Y, Zhang Y, Ouyang L. Targeting epigenetic modifications in Parkinson’s disease therapy. Medicinal Research Reviews. 2023;43(5):1748–1777. doi:10.1002/med.21962

[16] Bose A, Beal MF. Mitochondrial dysfunction in Parkinson’s disease. Journal of Neurochemistry. 2016;139(S1):216–231. doi:10.1111/jnc.13731

[17] Gao X-Y, Yang T, Gu Y, Sun X-H. Mitochondrial Dysfunction in Parkinson’s Disease: From Mechanistic Insights to Therapy. Frontiers in Aging Neuroscience. 2022;14:885500. doi:10.3389/fnagi.2022.885500

[18] Expanding the role of proteasome homeostasis in Parkinson’s disease: beyond protein breakdown | Cell Death & Disease. [accessed 2026 Oct 20]. https://www.nature.com/articles/s41419-021-03441-0

[19] Riederer P, Reichmann H, Youdim MBH, Gerlach M, editors. Parkinson’s Disease and Related Disorders. Vienna: Springer Vienna; 2006. http://link.springer.com/10.1007/978-3-211-45295-0. doi:10.1007/978-3-211-45295-0

[20] Sun X, Gu R, Bai J. Differentiation and regulation of CD4+ T cell subsets in Parkinson’s disease. Cellular and molecular life sciences: CMLS. 2024;81(1):352. doi:10.1007/s00018-024-05402-0

[21] Shebl N, Salama M. Exploring dopa decarboxylase as an ideal biomarker in Parkinson’s disease with focus on regulatory mechanisms, cofactor influences, and metabolic implications. npj Biomedical Innovations. 2025;2(1):2. doi:10.1038/s44385-024-00005-7

[22] Bolsewig K, Bellomo G, Hok-A-Hin YS, Al Idrissi I, Vermunt L, Lleó A, Alcolea D, Sieben A, Engelborghs S, Simonsen AH, et al. A quantitative DOPA decarboxylase biomarker for diagnosis in Lewy body disorders. Nature Medicine. 2026;32(3):1073–1084. doi:10.1038/s41591-026-04212-0

[23] Jenner P. Molecular mechanisms of L-DOPA-induced dyskinesia. Nature Reviews Neuroscience. 2008;9(9):665–677. doi:10.1038/nrn2471

[24] van Hillegondsberg L, Ahmad S, Zerenner T, Lawton M, Groenewald K, Fletcher M, Ianniello A, Ben-Shlomo Y, Wade-Martins R, Piazza P, et al. Dopamine replacement therapy modulates DDC, prolactin and AOC3 levels in Parkinson’s disease. npj Parkinson’s Disease. 2026;12(1):263. doi:10.1038/s41531-026-01435-3

[25] Vas S, Papp RS, Könczöl K, Bogáthy E, Papp N, Ádori C, Durst M, Sípos K, Ocskay K, Farkas I, et al. Prolactin-Releasing Peptide Contributes to Stress-Related Mood Disorders and Inhibits Sleep/Mood Regulatory Melanin-Concentrating Hormone Neurons in Rats. The Journal of Neuroscience. 2023;43(5):846. doi:10.1523/JNEUROSCI.2139-21.2022

[26] Fitzgerald P, Dinan TG. Prolactin and dopamine: What is the connection? A Review Article. Journal of Psychopharmacology. 2008;22(2_suppl):12–19. doi:10.1177/0269216307087148

[27] Scanga A, Lafontaine A-L, Kaminska M. An overview of the effects of levodopa and dopaminergic agonists on sleep disorders in Parkinson’s disease. Journal of Clinical Sleep Medicine. 2023;19(6):1133–1144. doi:10.5664/jcsm.10450

[28] Christine CW, Auinger P, Saleh N, Tian M, Bottiglieri T, Arning E, Tran NK, Ueland PM, Green R. Relationship of Cerebrospinal Fluid Vitamin B12 Status Markers With Parkinson’s Disease Progression. Movement Disorders. 2020;35(8):1466–1471. doi:10.1002/mds.28073

[29] Jia L, Wang Y, Wei W, Zhao W, Lu F, Liu F. Vitamin B12 inhibits α-synuclein fibrillogenesis and protects against amyloid-induced cytotoxicity. Food & Function. 2019;10(5):2861–2870. doi:10.1039/c8fo02471e

[30] Vijiaratnam N, Girges C, Auld G, McComish R, King A, Skene SS, Hibbert S, Wong A, Melander S, Gibson R, et al. Exenatide once a week versus placebo as a potential disease-modifying treatment for people with Parkinson’s disease in the UK: a phase 3, multicentre, double-blind, parallel-group, randomised, placebo-controlled trial. The Lancet. 2025;405(10479):627–636. doi:10.1016/S0140-6736(24)02808-3

[31] Curry DW, Stutz B, Andrews ZB, Elsworth JD. Targeting AMPK Signaling as a Neuroprotective Strategy in Parkinson’s Disease. Journal of Parkinson’s Disease. 2018;8(2):161–181. doi:10.3233/JPD-171296

[32] Szeliga M. Peroxiredoxins in Neurodegenerative Diseases. Antioxidants. 2020;9(12):1203. doi:10.3390/antiox9121203

[33] Love MI, Huber W, Anders S. Moderated estimation of fold change and dispersion for RNA-seq data with DESeq2. Genome Biology. 2014;15(12):550. doi:10.1186/s13059-014-0550-8

[34] Robinson MD, Oshlack A. A scaling normalization method for differential expression analysis of RNA-seq data. Genome Biology. 2010;11(3):R25. doi:10.1186/gb-2010-11-3-r25

[35] Thomas PD, Ebert D, Muruganujan A, Mushayahama T, Albou L-P, Mi H. PANTHER: Making genome-scale phylogenetics accessible to all. Protein Science. 2022;31(1):8–22. doi:10.1002/pro.4218

[36] Korotkevich G, Sukhov V, Budin N, Shpak B, Artyomov MN, Sergushichev A. Fast gene set enrichment analysis. 2021 [accessed 2026 Sept 18]:060012. https://www.biorxiv.org/content/10.1101/060012v3. doi:10.1101/060012

[37] Toro-Domínguez D, Martorell-Marugán J, Martinez-Bueno M, López-Domínguez R, Carnero-Montoro E, Barturen G, Goldman D, Petri M, Carmona-Sáez P, Alarcón-Riquelme ME. Scoring personalized molecular portraits identify Systemic Lupus Erythematosus subtypes and predict individualized drug responses, symptomatology and disease progression. Briefings in Bioinformatics. 2022;23(5):bbac332. doi:10.1093/bib/bbac332

[38] Ritchie ME, Phipson B, Wu D, Hu Y, Law CW, Shi W, Smyth GK. limma powers differential expression analyses for RNA-sequencing and microarray studies. Nucleic Acids Research. 2015;43(7):e47. doi:10.1093/nar/gkv007

[39] Smyth GK, Ritchie M, Thorne N, Wettenhall J, Shi W, Hu Y. Linear Models for Microarray and RNA-Seq Data User’s Guide.

[40] Vargas-Fernández M, Luna J de D, Martorell-Marugán J, Carmona-Sáez P. Network analysis of longitudinal electronic health records using linear mixed models. BioData Mining. 2026;19(1):7. doi:10.1186/s13040-025-00508-y

