## Supplementary for "Longitudinal network analysis links multi-omics data with clinical outcomes in Parkinson’s disease"

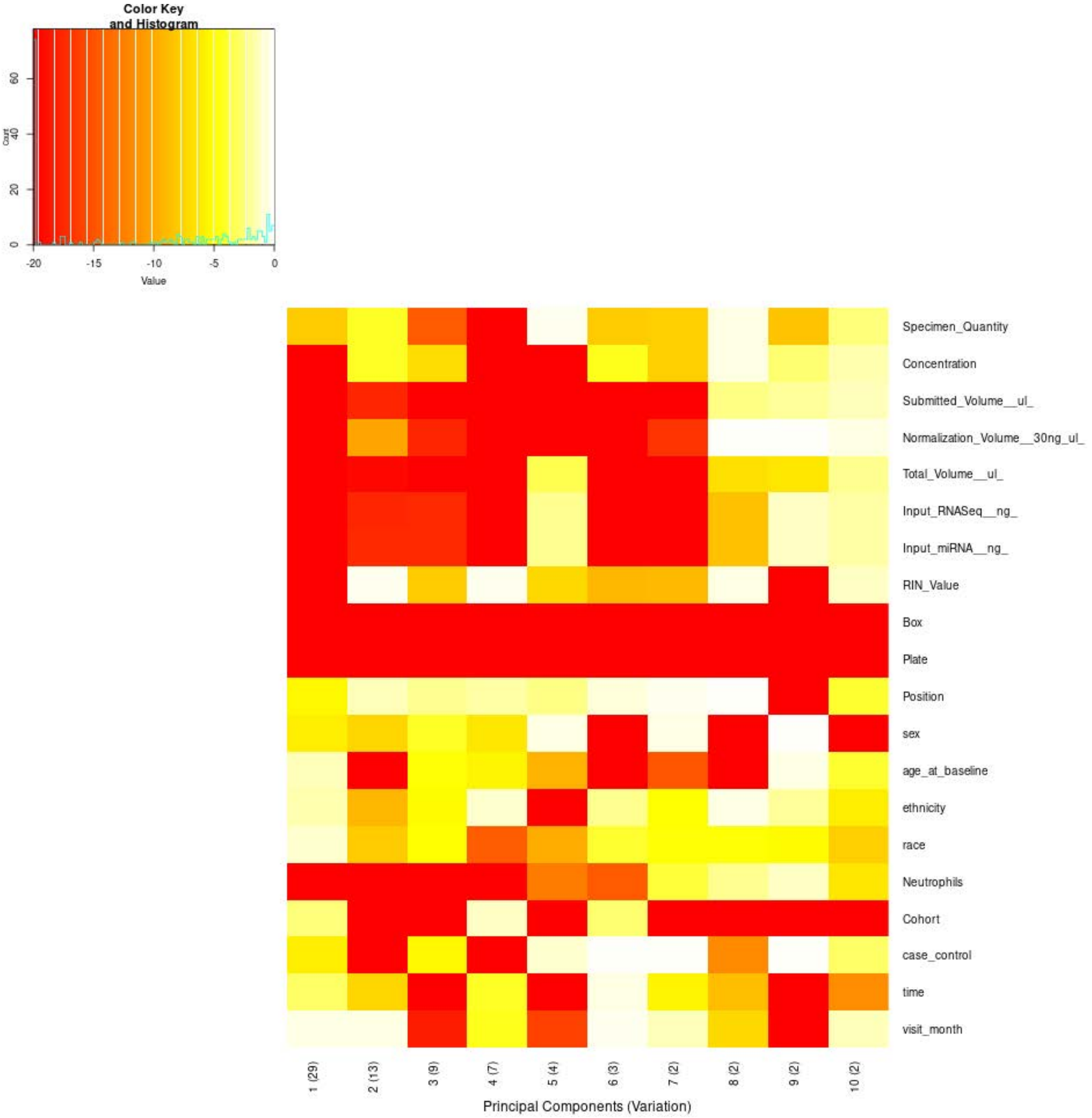


**Supplementary Figure 1.** Association of technical and clinical variables with the principal components of the whole-blood transcriptomics data before covariate correction (swamp prince plot). Each cell shows the log10 p-value of the association between a sample variable and one of the first ten principal components (percentage of variance explained in parentheses); red indicates strong association, white/pale yellow no association.

**Supplementary Table 1.** Values underlying Supplementary Figure 1 (whole-blood transcriptomics, before covariate correction). Top: p-values of the association between each sample variable and each of the first ten principal components (percentage of variance explained in parentheses). Bottom: corresponding $R^2$. $R^2 \geq 0.10$ in bold. <1E-300, below numerical precision.

| **Variable** | **PC1 (29%)** | **PC2 (13%)** | **PC3 (9%)** | **PC4 (7%)** | **PC5 (4%)** | **PC6 (3%)** | **PC7 (2%)** | **PC8 (2%)** | **PC9 (2%)** | **PC10 (2%)** |
|---|---|---|---|---|---|---|---|---|---|---|
| ***p-value*** | | | | | | | | | | |
| Specimen_Quantity | 9.81E-9 | 6.17E-5 | 2.55E-15 | 1.33E-36 | 5.56E-1 | 1.01E-8 | 1.40E-8 | 3.37E-1 | 3.42E-9 | 3.05E-3 |
| Concentration | 6.34E-47 | 6.07E-5 | 1.19E-7 | 9.92E-280 | 1.28E-20 | 3.60E-5 | 1.58E-8 | 3.77E-1 | 1.72E-3 | 2.98E-2 |
| Submitted_Volume__ul_ | 6.15E-81 | 1.82E-18 | 1.52E-30 | 2.44E-226 | 3.03E-33 | 1.21E-39 | 1.20E-32 | 3.98E-3 | 1.37E-2 | 4.29E-2 |
| Normalization_Volume__30ng_ul_ | 1.55E-83 | 5.82E-11 | 2.42E-18 | 3.88E-294 | 1.00E-57 | 8.94E-28 | 1.23E-17 | 6.57E-1 | 9.36E-1 | 2.58E-1 |
| Total_Volume__ul_ | 5.48E-40 | 3.95E-20 | 2.79E-32 | 2.86E-67 | 3.98E-4 | 3.39E-35 | 4.54E-38 | 1.59E-7 | 3.77E-7 | 8.66E-3 |
| Input_RNASeq__ng_ | 1.34E-23 | 2.65E-18 | 3.71E-18 | 1.03E-20 | 9.12E-3 | 4.29E-24 | 1.17E-33 | 2.93E-9 | 8.62E-2 | 2.06E-2 |
| Input_miRNA__ng_ | 1.34E-23 | 2.66E-18 | 3.73E-18 | 1.03E-20 | 9.11E-3 | 4.30E-24 | 1.18E-33 | 2.93E-9 | 8.63E-2 | 2.06E-2 |
| RIN_Value | 2.56E-113 | 6.09E-1 | 1.19E-8 | 5.22E-1 | 5.51E-8 | 6.50E-10 | 8.98E-10 | 3.12E-1 | 9.03E-83 | 6.62E-2 |
| Box | 5.94E-49 | 6.10E-73 | <1E-300 | 2.19E-27 | <1E-300 | 7.97E-227 | <1E-300 | 1.12E-52 | <1E-300 | 1.44E-154 |
| Plate | 7.92E-50 | 4.73E-75 | <1E-300 | 1.58E-28 | <1E-300 | 1.58E-237 | <1E-300 | 9.19E-54 | <1E-300 | 1.80E-158 |
| Position | 4.50E-6 | 4.59E-2 | 6.74E-3 | 1.94E-2 | 3.87E-3 | 1.76E-1 | 5.18E-1 | 7.74E-1 | 5.47E-26 | 1.02E-4 |
| sex | 1.11E-6 | 4.72E-8 | 6.89E-5 | 3.53E-7 | 3.02E-1 | 9.34E-26 | 3.65E-1 | 2.21E-165 | 7.62E-1 | 1.25E-53 |
| age_at_baseline | 5.70E-2 | 2.45E-34 | 1.52E-5 | 2.36E-6 | 3.61E-10 | 3.42E-41 | 1.32E-15 | 2.21E-49 | 2.99E-1 | 1.25E-4 |
| ethnicity | 2.59E-2 | 7.75E-10 | 7.07E-6 | 1.19E-1 | 8.61E-22 | 9.06E-3 | 1.16E-5 | 3.51E-1 | 1.22E-2 | 1.22E-6 |
| race | 1.09E-1 | 1.04E-8 | 1.03E-5 | 3.82E-15 | 1.66E-10 | 1.44E-4 | 1.53E-5 | 2.19E-5 | 6.73E-6 | 1.77E-8 |
| Neutrophils | 1.01E-89 | <1E-300 | 1.16E-173 | 3.79E-43 | 2.44E-13 | 1.90E-15 | 2.11E-4 | 7.71E-3 | 8.62E-2 | 3.94E-7 |
| Cohort | 2.54E-3 | 6.56E-69 | 4.96E-78 | 8.10E-2 | 7.60E-242 | 1.49E-3 | 4.85E-53 | 3.93E-22 | 6.71E-100 | 5.00E-86 |
| case_control | 1.38E-6 | 1.15E-24 | 3.92E-6 | 7.71E-28 | 1.35E-1 | 6.69E-1 | 6.67E-1 | 1.43E-12 | 9.85E-1 | 1.30E-3 |
| time | 1.09E-3 | 4.13E-8 | 6.11E-29 | 8.35E-5 | 9.28E-53 | 3.27E-1 | 3.19E-6 | 1.36E-9 | 4.07E-79 | 2.41E-12 |
| visit_month | 3.24E-1 | 3.59E-1 | 4.16E-19 | 4.59E-5 | 1.06E-16 | 4.94E-1 | 4.75E-2 | 6.44E-8 | 8.38E-32 | 5.08E-2 |
| ***$R^2$*** | | | | | | | | | | |
| Specimen_Quantity | 0.006 | 0.003 | 0.011 | 0.027 | 0.000 | 0.006 | 0.006 | 0.000 | 0.006 | 0.002 |
| Concentration | 0.035 | 0.003 | 0.005 | **0.200** | 0.015 | 0.003 | 0.006 | 0.000 | 0.002 | 0.001 |
| Submitted_Volume__ul_ | 0.061 | 0.013 | 0.023 | **0.165** | 0.025 | 0.030 | 0.024 | 0.001 | 0.001 | 0.001 |
| Normalization_Volume__30ng_ul_ | 0.063 | 0.007 | 0.013 | **0.209** | 0.044 | 0.021 | 0.013 | 0.000 | 0.000 | 0.000 |
| Total_Volume__ul_ | 0.030 | 0.015 | 0.024 | 0.051 | 0.002 | 0.026 | 0.029 | 0.005 | 0.004 | 0.001 |
| Input_RNASeq__ng_ | 0.017 | 0.013 | 0.013 | 0.015 | 0.001 | 0.018 | 0.025 | 0.006 | 0.001 | 0.001 |
| Input_miRNA__ng_ | 0.017 | 0.013 | 0.013 | 0.015 | 0.001 | 0.018 | 0.025 | 0.006 | 0.001 | 0.001 |
| RIN_Value | 0.085 | 0.000 | 0.006 | 0.000 | 0.005 | 0.007 | 0.007 | 0.000 | 0.063 | 0.001 |
| Box | 0.075 | 0.097 | **0.397** | 0.054 | **0.286** | **0.213** | **0.288** | 0.079 | **0.427** | **0.161** |
| Plate | 0.078 | **0.100** | **0.411** | 0.057 | **0.294** | **0.222** | **0.297** | 0.082 | **0.443** | **0.167** |
| Position | 0.029 | 0.020 | 0.023 | 0.021 | 0.023 | 0.018 | 0.016 | 0.014 | 0.054 | 0.026 |
| sex | 0.004 | 0.005 | 0.003 | 0.005 | 0.000 | 0.019 | 0.000 | **0.123** | 0.000 | 0.041 |
| age_at_baseline | 0.001 | 0.026 | 0.003 | 0.004 | 0.007 | 0.031 | 0.011 | 0.037 | 0.000 | 0.003 |
| ethnicity | 0.001 | 0.007 | 0.004 | 0.001 | 0.017 | 0.002 | 0.004 | 0.000 | 0.002 | 0.005 |
| race | 0.002 | 0.008 | 0.006 | 0.014 | 0.010 | 0.005 | 0.006 | 0.005 | 0.006 | 0.008 |
| Neutrophils | 0.068 | **0.635** | **0.129** | 0.033 | 0.009 | 0.011 | 0.002 | 0.001 | 0.001 | 0.004 |
| Cohort | 0.002 | 0.053 | 0.060 | 0.001 | **0.176** | 0.002 | 0.041 | 0.017 | 0.077 | 0.066 |
| case_control | 0.004 | 0.018 | 0.004 | 0.021 | 0.000 | 0.000 | 0.000 | 0.009 | 0.000 | 0.002 |
| time | 0.004 | 0.008 | 0.025 | 0.005 | 0.044 | 0.001 | 0.006 | 0.009 | 0.064 | 0.012 |
| visit_month | 0.000 | 0.000 | 0.014 | 0.003 | 0.012 | 0.000 | 0.001 | 0.005 | 0.024 | 0.001 |

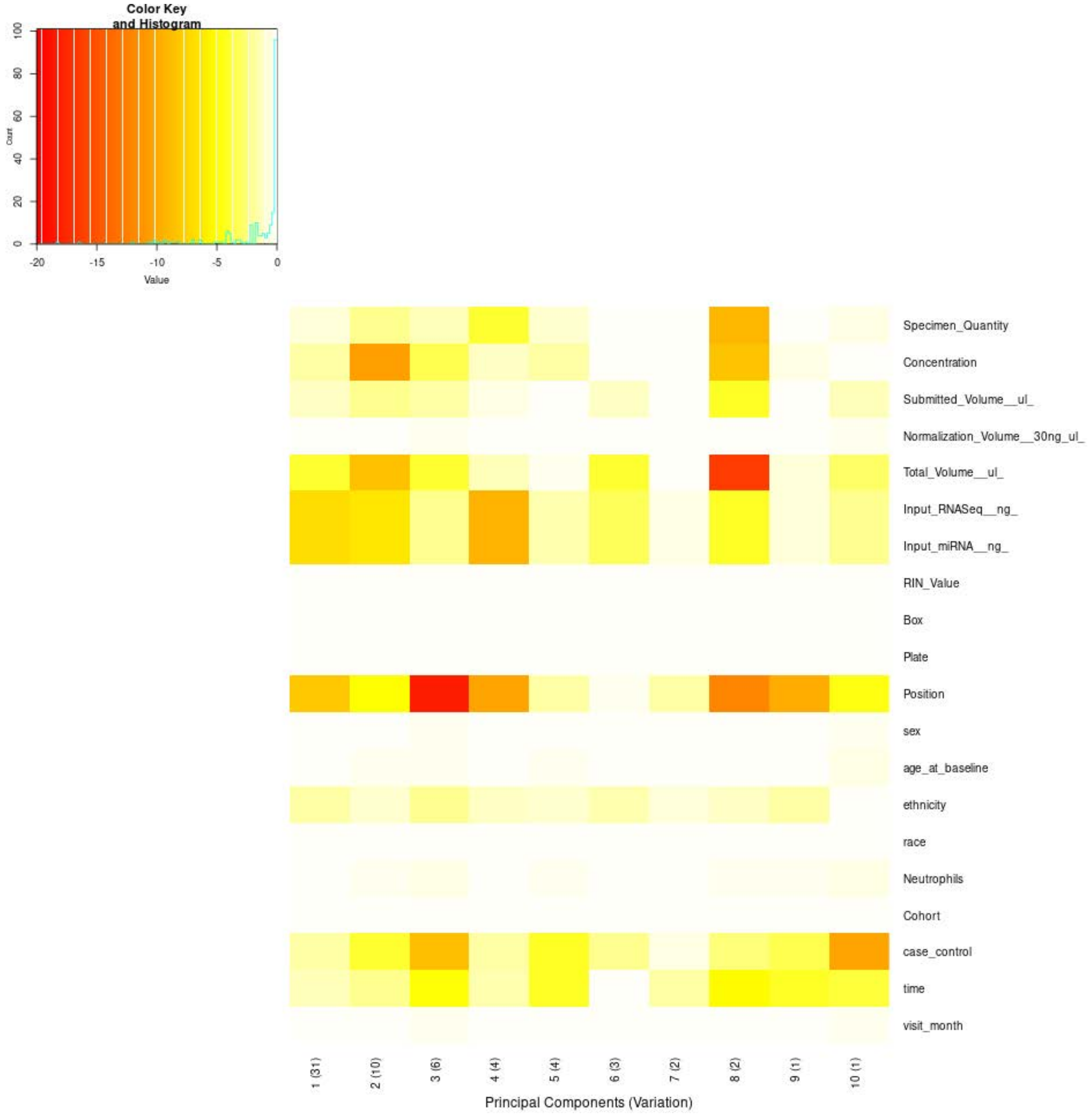


**Supplementary Figure 2.** Association of technical and clinical variables with the principal components of the whole-blood transcriptomics data after covariate correction (swamp prince plot). Each cell shows the log10 p-value of the association between a sample variable and one of the first ten principal components (percentage of variance explained in parentheses); red indicates strong association, white/pale yellow no association.

**Supplementary Table 2.** Values underlying Supplementary Figure 2 (whole-blood transcriptomics, after covariate correction). Top: p-values of the association between each sample variable and each of the first ten principal components (percentage of variance explained in parentheses). Bottom: corresponding $R^2$. $R^2 \geq 0.10$ in bold. <1E-300, below numerical precision.

| Variable | PC1 (31%) | PC2 (10%) | PC3 (6%) | PC4 (4%) | PC5 (4%) | PC6 (3%) | PC7 (2%) | PC8 (2%) | PC9 (1%) | PC10 (1%) |
|---|---|---|---|---|---|---|---|---|---|---|
| ***p-value*** | | | | | | | | | | |
| Specimen_Quantity | 1.60E-1 | 8.31E-3 | 5.70E-2 | 1.05E-4 | 1.00E-1 | 8.84E-1 | 7.06E-1 | 5.49E-10 | 9.68E-1 | 3.19E-1 |
| Concentration | 1.73E-2 | 3.07E-11 | 4.40E-4 | 7.03E-2 | 1.52E-2 | 9.07E-1 | 9.46E-1 | 4.47E-9 | 2.63E-1 | 9.84E-1 |
| Submitted_Volume__ul_ | 7.72E-2 | 6.72E-3 | 2.33E-2 | 3.89E-1 | 9.30E-1 | 8.25E-2 | 9.70E-1 | 6.16E-5 | 7.12E-1 | 5.00E-2 |
| Normalization_Volume__30ng_ul_ | 8.36E-1 | 7.24E-1 | 5.85E-1 | 8.33E-1 | 7.18E-1 | 8.05E-1 | 9.17E-1 | 7.92E-1 | 7.52E-1 | 5.53E-1 |
| Total_Volume__ul_ | 1.25E-4 | 2.87E-9 | 1.09E-4 | 4.49E-2 | 5.16E-1 | 1.28E-4 | 9.52E-1 | 2.81E-17 | 2.48E-1 | 1.45E-3 |
| Input_RNASeq__ng_ | 1.04E-7 | 4.48E-7 | 7.19E-3 | 4.78E-10 | 2.59E-2 | 6.79E-4 | 3.53E-1 | 6.31E-5 | 1.86E-1 | 8.05E-3 |
| Input_miRNA__ng_ | 1.04E-7 | 4.48E-7 | 7.19E-3 | 4.78E-10 | 2.58E-2 | 6.79E-4 | 3.53E-1 | 6.31E-5 | 1.86E-1 | 8.05E-3 |
| RIN_Value | 9.01E-1 | 8.32E-1 | 7.43E-1 | 8.99E-1 | 8.28E-1 | 8.82E-1 | 9.50E-1 | 8.74E-1 | 8.49E-1 | 7.22E-1 |
| Box | 1.00E0 | 1.00E0 | 1.00E0 | 1.00E0 | 1.00E0 | 1.00E0 | 1.00E0 | 1.00E0 | 1.00E0 | 1.00E0 |
| Plate | 1.00E0 | 1.00E0 | 1.00E0 | 1.00E0 | 1.00E0 | 1.00E0 | 1.00E0 | 1.00E0 | 1.00E0 | 1.00E0 |
| Position | 8.22E-9 | 1.25E-5 | 5.90E-19 | 5.18E-11 | 2.40E-2 | 6.07E-1 | 2.17E-2 | 8.86E-13 | 1.54E-10 | 2.69E-5 |
| sex | 8.28E-1 | 7.11E-1 | 5.66E-1 | 8.25E-1 | 7.04E-1 | 7.95E-1 | 9.13E-1 | 7.82E-1 | 7.40E-1 | 5.33E-1 |
| age_at_baseline | 7.57E-1 | 5.98E-1 | 4.14E-1 | 7.53E-1 | 5.89E-1 | 7.12E-1 | 8.76E-1 | 6.94E-1 | 6.37E-1 | 3.75E-1 |
| ethnicity | 1.90E-2 | 9.99E-2 | 9.09E-3 | 8.32E-2 | 1.22E-1 | 3.68E-2 | 2.29E-1 | 8.24E-2 | 2.12E-2 | 9.62E-1 |
| race | 1.00E0 | 1.00E0 | 9.98E-1 | 1.00E0 | 1.00E0 | 1.00E0 | 1.00E0 | 1.00E0 | 1.00E0 | 9.97E-1 |
| Neutrophils | 6.95E-1 | 5.04E-1 | 3.01E-1 | 6.90E-1 | 4.94E-1 | 6.40E-1 | 8.44E-1 | 6.18E-1 | 5.50E-1 | 2.61E-1 |
| Cohort | 9.96E-1 | 9.89E-1 | 9.75E-1 | 9.96E-1 | 9.89E-1 | 9.95E-1 | 9.99E-1 | 9.94E-1 | 9.91E-1 | 9.70E-1 |
| case_control | 2.25E-2 | 9.83E-5 | 1.55E-9 | 2.01E-2 | 6.65E-5 | 6.49E-3 | 2.51E-1 | 3.62E-3 | 4.86E-4 | 5.56E-11 |
| time | 5.15E-2 | 7.94E-3 | 1.85E-5 | 3.84E-2 | 5.86E-5 | 9.53E-1 | 2.14E-2 | 8.44E-6 | 6.91E-5 | 2.24E-4 |
| visit_month | 8.38E-1 | 7.27E-1 | 5.89E-1 | 8.35E-1 | 7.21E-1 | 8.07E-1 | 9.18E-1 | 7.94E-1 | 7.55E-1 | 5.58E-1 |
| ***$R^2$*** | | | | | | | | | | |
| Specimen_Quantity | 0.000 | 0.001 | 0.001 | 0.003 | 0.000 | 0.000 | 0.000 | 0.007 | 0.000 | 0.000 |
| Concentration | 0.001 | 0.008 | 0.002 | 0.001 | 0.001 | 0.000 | 0.000 | 0.006 | 0.000 | 0.000 |
| Submitted_Volume__ul_ | 0.001 | 0.001 | 0.001 | 0.000 | 0.000 | 0.001 | 0.000 | 0.003 | 0.000 | 0.001 |
| Normalization_Volume__30ng_ul_ | 0.000 | 0.000 | 0.000 | 0.000 | 0.000 | 0.000 | 0.000 | 0.000 | 0.000 | 0.000 |
| Total_Volume__ul_ | 0.003 | 0.006 | 0.003 | 0.001 | 0.000 | 0.003 | 0.000 | 0.012 | 0.000 | 0.002 |
| Input_RNASeq__ng_ | 0.005 | 0.004 | 0.001 | 0.007 | 0.001 | 0.002 | 0.000 | 0.003 | 0.000 | 0.001 |
| Input_miRNA__ng_ | 0.005 | 0.004 | 0.001 | 0.007 | 0.001 | 0.002 | 0.000 | 0.003 | 0.000 | 0.001 |
| RIN_Value | 0.000 | 0.000 | 0.000 | 0.000 | 0.000 | 0.000 | 0.000 | 0.000 | 0.000 | 0.000 |
| Box | 0.000 | 0.001 | 0.001 | 0.001 | 0.001 | 0.000 | 0.001 | 0.001 | 0.001 | 0.000 |
| Plate | 0.000 | 0.000 | 0.000 | 0.000 | 0.000 | 0.000 | 0.000 | 0.000 | 0.000 | 0.000 |
| Position | 0.033 | 0.028 | 0.046 | 0.036 | 0.021 | 0.015 | 0.021 | 0.039 | 0.036 | 0.028 |
| sex | 0.000 | 0.000 | 0.000 | 0.000 | 0.000 | 0.000 | 0.000 | 0.000 | 0.000 | 0.000 |
| age_at_baseline | 0.000 | 0.000 | 0.000 | 0.000 | 0.000 | 0.000 | 0.000 | 0.000 | 0.000 | 0.000 |
| ethnicity | 0.001 | 0.001 | 0.002 | 0.001 | 0.001 | 0.001 | 0.001 | 0.001 | 0.001 | 0.000 |
| race | 0.000 | 0.000 | 0.000 | 0.000 | 0.000 | 0.000 | 0.000 | 0.000 | 0.000 | 0.000 |
| Neutrophils | 0.000 | 0.000 | 0.000 | 0.000 | 0.000 | 0.000 | 0.000 | 0.000 | 0.000 | 0.000 |
| Cohort | 0.000 | 0.000 | 0.000 | 0.000 | 0.000 | 0.000 | 0.000 | 0.000 | 0.000 | 0.000 |
| case_control | 0.001 | 0.003 | 0.006 | 0.001 | 0.003 | 0.001 | 0.000 | 0.001 | 0.002 | 0.007 |
| time | 0.002 | 0.003 | 0.006 | 0.002 | 0.005 | 0.000 | 0.003 | 0.006 | 0.005 | 0.005 |
| visit_month | 0.000 | 0.000 | 0.000 | 0.000 | 0.000 | 0.000 | 0.000 | 0.000 | 0.000 | 0.000 |

**Supplementary Table 3.** Selection of covariates for the whole-blood transcriptomics differential expression model. Variance explained is the $R^2$ of the association between each sample variable and each of the first ten principal components, weighted by the proportion of variance explained by that component, before and after covariate correction (Supplementary Figures 1 and 2; Supplementary Tables 1 and 2). Sig. PCs, number of principal components significantly associated with the variable (FDR < 0.05, Benjamini–Hochberg correction within each component). Model: Covariate, included in the model; Diagnosis, variable of interest, preserved during correction; —, not included. Top correlated covariate, numeric sample variable with the highest absolute Spearman correlation (ρ); not applicable to categorical variables (—). Within blocks of strongly correlated technical variables (|ρ| ≥ 0.99), the variable explaining the most variance was retained. Box is equivalent to sequencing plate, and plates are nested within cohort; time is the visit code underlying visit_month. Variables are ordered by variance explained before correction.

| Variable | Model | Variance explained before (%) | Sig. PCs before | Variance explained after (%) | Sig. PCs after | Top correlated covariate (ρ) |
|---|---|---|---|---|---|---|
| Neutrophils | Covariate | 11.73 | 9 | 0.00 | 0 | age_at_baseline (0.109) |
| Plate | Covariate | 11.21 | 10 | 0.00 | 0 | — |
| Box | — | 10.81 | 10 | 0.04 | 0 | — |
| Normalization_Volume__30ng_ul_ | Covariate | 3.69 | 7 | 0.00 | 0 | Submitted_Volume__ul_ (-0.990) |
| Submitted_Volume__ul_ | — | 3.48 | 9 | 0.04 | 2 | Concentration (-0.996) |
| RIN_Value | Covariate | 2.66 | 6 | 0.00 | 0 | Normalization_Volume__30ng_ul_ (-0.164) |
| Concentration | — | 2.52 | 9 | 0.14 | 3 | Submitted_Volume__ul_ (-0.996) |
| Cohort | — | 2.36 | 9 | 0.00 | 0 | — |
| Position | — | 1.78 | 6 | 2.10 | 7 | — |
| Total_Volume__ul_ | — | 1.78 | 10 | 0.20 | 6 | Input_RNASeq__ng_ (-0.640) |
| Input_miRNA__ng_ | — | 1.02 | 9 | 0.25 | 7 | Input_RNASeq__ng_ (1.000) |
| Input_RNASeq__ng_ | — | 1.02 | 9 | 0.25 | 7 | Input_miRNA__ng_ (1.000) |
| time | — | 0.81 | 9 | 0.19 | 6 | — |
| age_at_baseline | Covariate | 0.63 | 8 | 0.00 | 0 | Neutrophils (0.109) |
| sex | Covariate | 0.62 | 7 | 0.00 | 0 | — |
| case_control | Diagnosis | 0.55 | 6 | 0.13 | 7 | — |
| Specimen_Quantity | — | 0.52 | 8 | 0.05 | 3 | Input_RNASeq__ng_ (0.597) |
| race | Covariate | 0.41 | 9 | 0.00 | 0 | — |
| ethnicity | — | 0.26 | 8 | 0.08 | 1 | — |
| visit_month | Covariate | 0.25 | 5 | 0.00 | 0 | Specimen_Quantity (-0.123) |

**Supplementary Table 4.** Differentially expressed genes between PD cases and controls in whole-blood transcriptomics data. Genes with padj < 0.05 and |log2FoldChange| > 0.15 are shown, ordered by absolute log2 fold change; positive log2FoldChange values indicate higher expression in PD cases. baseMean, mean of normalized counts across samples; log2FoldChange, log2 fold change; lfcSE, standard error of the log2 fold change; stat, Wald statistic; pvalue, Wald test p-value; padj, Benjamini–Hochberg adjusted p-value; abslogFC, absolute log2 fold change.

| Gene | baseMean | log2FoldChange | lfcSE | stat | pvalue | padj | abslogFC |
|---|---|---|---|---|---|---|---|
| *FOLR3* | 752.86782 | 0.19655 | 0.02969 | 6.62079 | 3.57275E-11 | 1.70063E-8 | 0.19655 |
| *H1-0* | 923.81949 | -0.18076 | 0.02414 | -7.48660 | 7.06801E-14 | 5.71944E-11 | 0.18076 |
| *BPGM* | 3694.92019 | -0.17698 | 0.02322 | -7.62141 | 2.50914E-14 | 3.82510E-11 | 0.17698 |
| *XK* | 1786.70441 | -0.17639 | 0.02334 | -7.55661 | 4.13724E-14 | 4.37902E-11 | 0.17639 |
| *CA1* | 2644.93605 | -0.17466 | 0.02971 | -5.87910 | 4.12507E-9 | 6.67601E-7 | 0.17466 |
| *PITHD1* | 4589.59760 | -0.16463 | 0.02180 | -7.55070 | 4.32924E-14 | 4.37902E-11 | 0.16463 |
| *FECH* | 5017.23901 | -0.16332 | 0.02147 | -7.60558 | 2.83621E-14 | 3.82510E-11 | 0.16332 |
| *RN7SL3* | 42532.10529 | -0.16258 | 0.02953 | -5.50649 | 3.66053E-8 | 4.11403E-6 | 0.16258 |
| *SLC6A8* | 911.05368 | -0.15731 | 0.02542 | -6.18898 | 6.05565E-10 | 1.37090E-7 | 0.15731 |
| *ANKRD36BP2* | 812.73635 | -0.15482 | 0.02809 | -5.51163 | 3.55526E-8 | 4.05199E-6 | 0.15482 |
| *SPTB* | 1646.07307 | -0.15424 | 0.02668 | -5.78107 | 7.42280E-9 | 1.05378E-6 | 0.15424 |
| *TENT5C* | 23869.17206 | -0.15412 | 0.02352 | -6.55357 | 5.61791E-11 | 2.39264E-8 | 0.15412 |
| *TRIM58* | 14201.67323 | -0.15068 | 0.02350 | -6.41142 | 1.44172E-10 | 4.86099E-8 | 0.15068 |

**Supplementary Table 5.** Gene Ontology Biological Process terms enriched in gene set enrichment analysis (fgsea) of the ranked transcriptomic differential expression results. Terms with padj < 0.05 and |NES| ≥ 1.5 are shown, ordered by padj. Positive NES indicates enrichment among genes with higher expression in PD cases. pval, nominal p-value; padj, Benjamini–Hochberg adjusted p-value; log2err, expected error of the log2 p-value estimate; ES, enrichment score; NES, normalized enrichment score; size, number of genes in the term after filtering; Collapsed into, for terms identified as redundant by collapsePathways (fgsea), the term that accounts for their enrichment; —, non-redundant term (13 terms) used in downstream analyses.

| Pathway | Term | pval | padj | log2err | ES | NES | size | Collapsed into |
|---|---|---|---|---|---|---|---|---|
| GO:0006119 | oxidative phosphorylation | 3.35802E-6 | 0.00293 | 0.62726 | -0.82431 | -2.25954 | 16 | — |
| GO:0007165 | signal transduction | 2.41375E-6 | 0.00293 | 0.62726 | 0.28919 | 1.52870 | 586 | — |
| GO:0007186 | G protein-coupled receptor signaling pathway | 4.06778E-6 | 0.00293 | 0.61053 | 0.43783 | 1.95490 | 126 | — |
| GO:0045653 | negative regulation of megakaryocyte differentiation | 3.89055E-6 | 0.00293 | 0.61053 | -0.84888 | -2.16737 | 13 | — |
| GO:0035556 | intracellular signal transduction | 7.80774E-6 | 0.00450 | 0.59333 | 0.35569 | 1.73417 | 231 | — |
| GO:0000209 | protein polyubiquitination | 1.53538E-5 | 0.00638 | 0.57561 | -0.44782 | -1.85708 | 138 | — |
| GO:0007200 | phospholipase C-activating G protein-coupled receptor signaling pathway | 1.55133E-5 | 0.00638 | 0.57561 | 0.81230 | 2.09800 | 12 | — |
| GO:0006335 | DNA replication-dependent chromatin assembly | 4.79797E-5 | 0.01727 | 0.55733 | -0.77670 | -2.12905 | 16 | — |
| GO:2000556 | positive regulation of T-helper 1 cell cytokine production | 7.66569E-5 | 0.02452 | 0.53843 | 0.92357 | 1.95365 | 6 | — |
| GO:0043161 | proteasome-mediated ubiquitin-dependent protein catabolic process | 8.91691E-5 | 0.02567 | 0.53843 | -0.42376 | -1.75889 | 141 | — |
| GO:0032200 | telomere organization | 1.19434E-4 | 0.03126 | 0.53843 | -0.76861 | -2.01451 | 14 | negative regulation of megakaryocyte differentiation |
| GO:0006511 | ubiquitin-dependent protein catabolic process | 1.43434E-4 | 0.03441 | 0.51885 | -0.37347 | -1.62245 | 198 | protein polyubiquitination |
| GO:0016567 | protein ubiquitination | 1.72469E-4 | 0.03820 | 0.51885 | -0.33290 | -1.54147 | 380 | — |
| GO:0006935 | chemotaxis | 1.85817E-4 | 0.03821 | 0.51885 | 0.49748 | 1.94808 | 59 | — |
| GO:0006325 | chromatin organization | 2.31202E-4 | 0.04438 | 0.51885 | 0.31640 | 1.54045 | 245 | — |
| GO:0006334 | nucleosome assembly | 2.76205E-4 | 0.04970 | 0.49849 | -0.51624 | -1.88486 | 61 | DNA replication-dependent chromatin assembly |

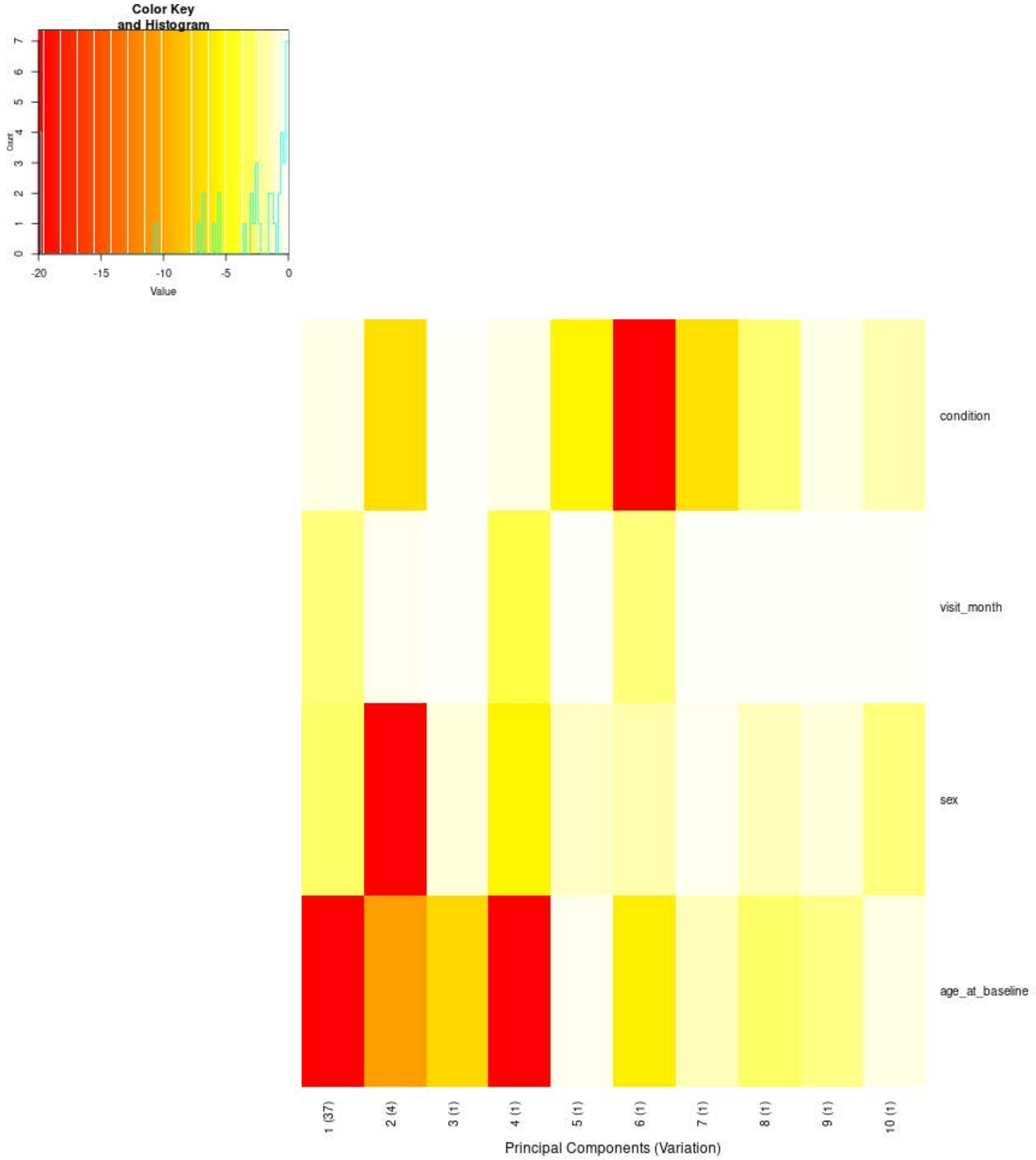


**Supplementary Figure 3.** Association of clinical variables with the principal components of the CSF proteomics data before covariate correction (swamp prince plot). Each cell shows the log10 p-value of the association between a sample variable and one of the first ten principal components (percentage of variance explained in parentheses); red indicates strong association, white/pale yellow no association.

**Supplementary Table 6.** Values underlying Supplementary Figure 3 (CSF proteomics, before covariate correction). Top: p-values of the association between each sample variable and each of the first ten principal components (percentage of variance explained in parentheses). Bottom: corresponding $R^2$. $R^2 \geq 0.10$ in bold.

| Variable | PC1 (37%) | PC2 (4%) | PC3 (1%) | PC4 (1%) | PC5 (1%) | PC6 (1%) | PC7 (1%) | PC8 (1%) | PC9 (1%) | PC10 (1%) |
|---|---|---|---|---|---|---|---|---|---|---|
| ***p-value*** | | | | | | | | | | |
| condition | 3.12E-1 | 1.48E-7 | 9.63E-1 | 3.64E-1 | 3.16E-6 | 5.72E-24 | 1.42E-7 | 2.14E-3 | 3.12E-1 | 3.73E-2 |
| visit_month | 3.49E-3 | 6.00E-1 | 6.99E-1 | 2.92E-4 | 7.98E-1 | 2.75E-3 | 7.60E-1 | 9.50E-1 | 7.57E-1 | 6.59E-1 |
| sex | 1.35E-3 | 7.35E-30 | 2.03E-1 | 3.07E-6 | 7.51E-2 | 3.66E-2 | 5.25E-1 | 4.38E-2 | 2.16E-1 | 3.32E-3 |
| age_at_baseline | 1.10E-22 | 2.35E-11 | 5.56E-8 | 5.45E-88 | 5.25E-1 | 1.20E-6 | 4.07E-2 | 1.46E-3 | 4.12E-3 | 3.81E-1 |
| ***$R^2$*** | | | | | | | | | | |
| condition | 0.002 | 0.024 | 0.000 | 0.002 | 0.019 | 0.078 | 0.024 | 0.009 | 0.002 | 0.005 |
| visit_month | 0.006 | 0.000 | 0.000 | 0.010 | 0.000 | 0.007 | 0.000 | 0.000 | 0.000 | 0.000 |
| sex | 0.008 | 0.093 | 0.001 | 0.016 | 0.002 | 0.003 | 0.000 | 0.003 | 0.001 | 0.007 |
| age_at_baseline | 0.070 | 0.033 | 0.022 | **0.259** | 0.000 | 0.018 | 0.003 | 0.008 | 0.006 | 0.001 |

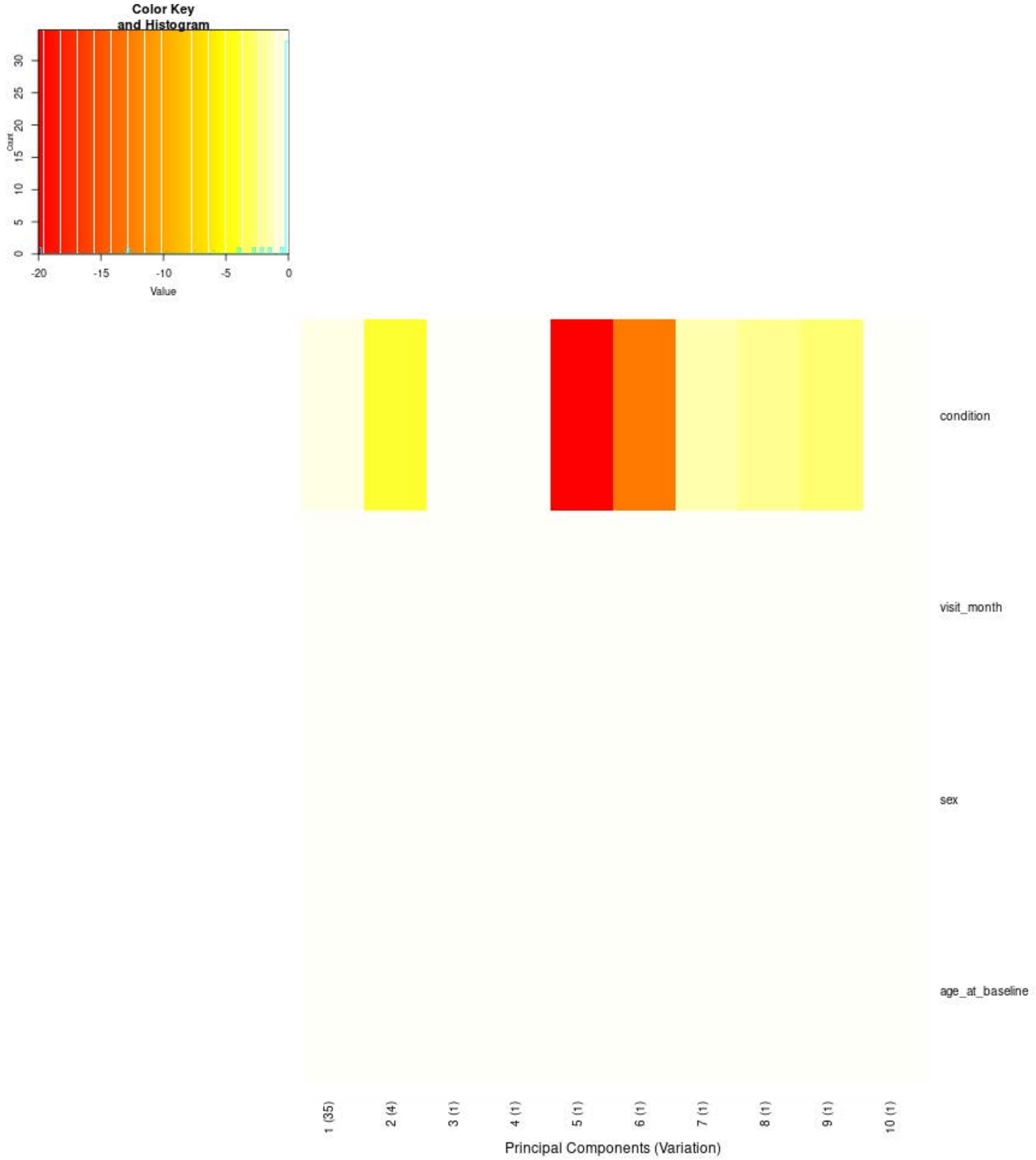


**Supplementary Figure 4.** Association of clinical variables with the principal components of the CSF proteomics data after correction for visit month, sex and age at baseline (swamp prince plot). Each cell shows the log10 p-value of the association between a sample variable and one of the first ten principal components (percentage of variance explained in parentheses); red indicates strong association, white/pale yellow no association.

**Supplementary Table 7.** Values underlying Supplementary Figure 4 (CSF proteomics, after correction for visit month, sex and age at baseline). Top: p-values of the association between each sample variable and each of the first ten principal components (percentage of variance explained in parentheses). Bottom: corresponding $R^2$. $R^2 \geq 0.10$ in bold.

| **Variable** | **PC1 (35%)** | **PC2 (4%)** | **PC3 (1%)** | **PC4 (1%)** | **PC5 (1%)** | **PC6 (1%)** | **PC7 (1%)** | **PC8 (1%)** | **PC9 (1%)** | **PC10 (1%)** |
|---|---|---|---|---|---|---|---|---|---|---|
| ***p-value*** | | | | | | | | | | |
| condition | 2.76E-1 | 1.34E-4 | 9.06E-1 | 7.24E-1 | 2.32E-24 | 1.49E-13 | 2.72E-2 | 7.14E-3 | 1.83E-3 | 8.84E-1 |
| visit_month | 1.00 | 1.00 | 1.00 | 1.00 | 1.00 | 1.00 | 1.00 | 1.00 | 1.00 | 1.00 |
| sex | 1.00 | 1.00 | 1.00 | 1.00 | 1.00 | 1.00 | 1.00 | 1.00 | 1.00 | 1.00 |
| age_at_baseline | 1.00 | 1.00 | 1.00 | 1.00 | 1.00 | 1.00 | 1.00 | 1.00 | 1.00 | 1.00 |
| ***R²*** | | | | | | | | | | |
| condition | 0.002 | 0.013 | 0.000 | 0.000 | 0.079 | 0.044 | 0.005 | 0.007 | 0.010 | 0.000 |
| visit_month | 0.000 | 0.000 | 0.000 | 0.000 | 0.000 | 0.000 | 0.000 | 0.000 | 0.000 | 0.000 |
| sex | 0.000 | 0.000 | 0.000 | 0.000 | 0.000 | 0.000 | 0.000 | 0.000 | 0.000 | 0.000 |
| age_at_baseline | 0.000 | 0.000 | 0.000 | 0.000 | 0.000 | 0.000 | 0.000 | 0.000 | 0.000 | 0.000 |

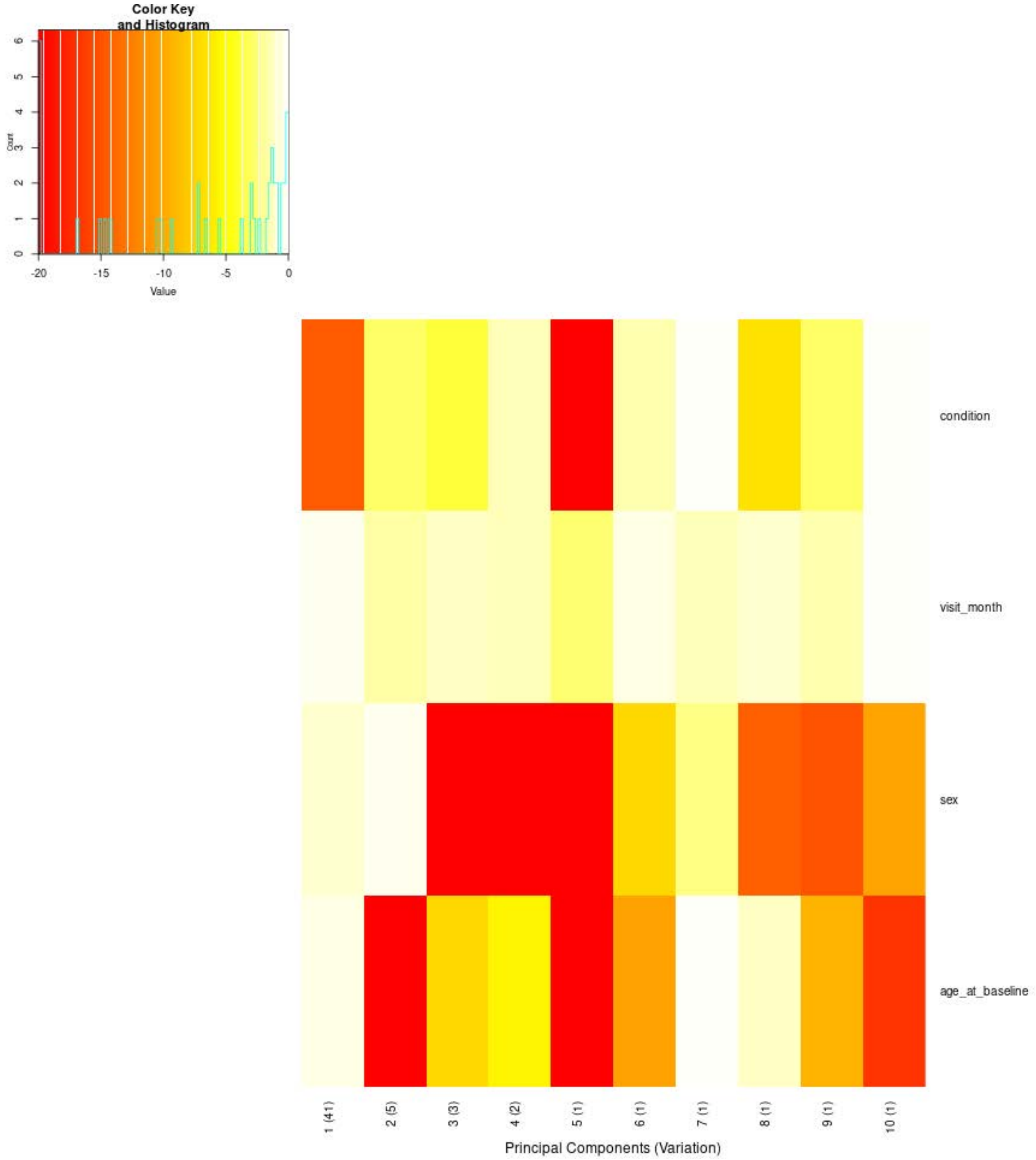


**Supplementary Figure 5.** Association of clinical variables with the principal components of the plasma proteomics data before covariate correction (swamp prince plot). Each cell shows the log10 p-value of the association between a sample variable and one of the first ten principal components (percentage of variance explained in parentheses); red indicates strong association, white/pale yellow no association.

**Supplementary Table 8.** Values underlying Supplementary Figure 5 (plasma proteomics, before covariate correction). Top: p-values of the association between each sample variable and each of the first ten principal components (percentage of variance explained in parentheses). Bottom: corresponding $R^2$. $R^2 \geq 0.10$ in bold.

| **Variable** | **PC1 (41%)** | **PC2 (5%)** | **PC3 (3%)** | **PC4 (2%)** | **PC5 (1%)** | **PC6 (1%)** | **PC7 (1%)** | **PC8 (1%)** | **PC9 (1%)** | **PC10 (1%)** |
|---|---|---|---|---|---|---|---|---|---|---|
| ***p-value*** | | | | | | | | | | |
| condition | 2.37E-15 | 1.23E-3 | 2.19E-4 | 5.22E-2 | 8.62E-26 | 2.56E-2 | 9.85E-1 | 2.92E-7 | 9.81E-4 | 8.89E-1 |
| visit_month | 4.17E-1 | 1.89E-2 | 9.34E-2 | 3.99E-2 | 1.59E-3 | 3.35E-1 | 5.65E-2 | 1.49E-1 | 3.79E-2 | 9.27E-1 |
| sex | 9.88E-2 | 5.79E-1 | 5.07E-28 | 1.38E-57 | 9.29E-36 | 8.39E-8 | 4.07E-3 | 5.18E-15 | 1.01E-15 | 5.68E-11 |
| age_at_baseline | 3.09E-1 | 2.55E-47 | 8.41E-8 | 3.35E-6 | 4.28E-29 | 3.36E-11 | 6.72E-1 | 6.91E-2 | 4.25E-10 | 1.12E-17 |
| ***$R^2$*** | | | | | | | | | | |
| condition | 0.043 | 0.009 | 0.011 | 0.004 | 0.072 | 0.005 | 0.000 | 0.019 | 0.009 | 0.000 |
| visit_month | 0.000 | 0.004 | 0.002 | 0.003 | 0.006 | 0.001 | 0.002 | 0.001 | 0.003 | 0.000 |
| sex | 0.002 | 0.000 | 0.075 | **0.152** | 0.096 | 0.018 | 0.005 | 0.039 | 0.041 | 0.027 |
| age_at_baseline | 0.001 | **0.126** | 0.018 | 0.014 | 0.078 | 0.028 | 0.000 | 0.002 | 0.025 | 0.046 |

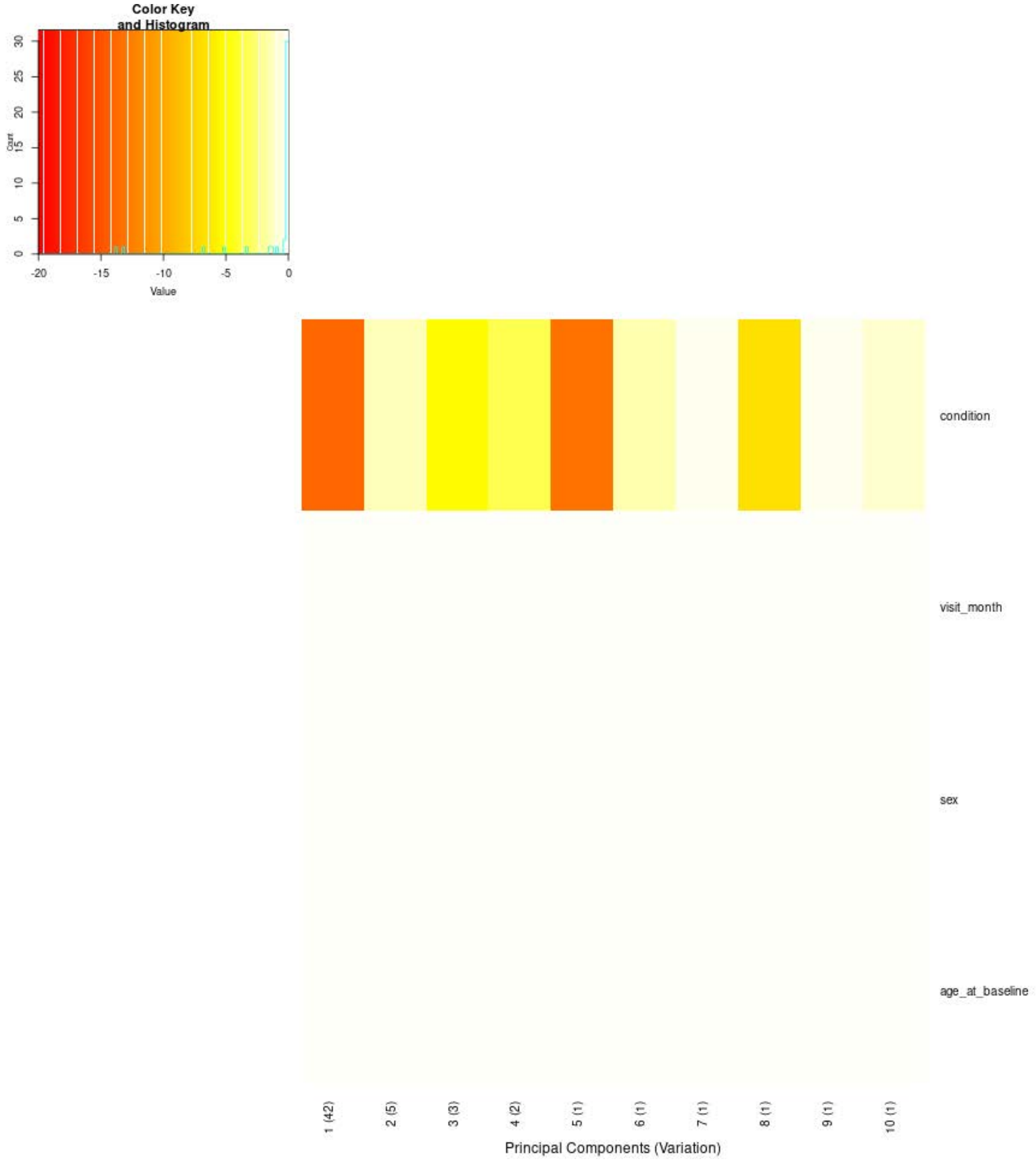


**Supplementary Figure 6.** Association of clinical variables with the principal components of the plasma proteomics data after correction for visit month, sex and age at baseline (swamp prince plot). Each cell shows the log10 p-value of the association between a sample variable and one of the first ten principal components (percentage of variance explained in parentheses); red indicates strong association, white/pale yellow no association.

**Supplementary Table 9.** Values underlying Supplementary Figure 6 (plasma proteomics, after correction for visit month, sex and age at baseline). Top: p-values of the association between each sample variable and each of the first ten principal components (percentage of variance explained in parentheses). Bottom: corresponding $R^2$. $R^2 \geq 0.10$ in bold.

| **Variable** | **PC1 (42%)** | **PC2 (5%)** | **PC3 (3%)** | **PC4 (2%)** | **PC5 (1%)** | **PC6 (1%)** | **PC7 (1%)** | **PC8 (1%)** | **PC9 (1%)** | **PC10 (1%)** |
|---|---|---|---|---|---|---|---|---|---|---|
| ***p-value*** | | | | | | | | | | |
| condition | 1.28E-14 | 5.52E-2 | 7.04E-6 | 4.98E-4 | 6.17E-14 | 3.65E-2 | 5.10E-1 | 1.48E-7 | 5.77E-1 | 1.06E-1 |
| visit_month | 1.00 | 1.00 | 1.00 | 1.00 | 1.00 | 1.00 | 1.00 | 1.00 | 1.00 | 1.00 |
| sex | 1.00 | 1.00 | 1.00 | 1.00 | 1.00 | 1.00 | 1.00 | 1.00 | 1.00 | 1.00 |
| age_at_baseline | 1.00 | 1.00 | 1.00 | 1.00 | 1.00 | 1.00 | 1.00 | 1.00 | 1.00 | 1.00 |
| ***$R^2$*** | | | | | | | | | | |
| condition | 0.040 | 0.004 | 0.015 | 0.010 | 0.039 | 0.004 | 0.001 | 0.020 | 0.001 | 0.003 |
| visit_month | 0.000 | 0.000 | 0.000 | 0.000 | 0.000 | 0.000 | 0.000 | 0.000 | 0.000 | 0.000 |
| sex | 0.000 | 0.000 | 0.000 | 0.000 | 0.000 | 0.000 | 0.000 | 0.000 | 0.000 | 0.000 |
| age_at_baseline | 0.000 | 0.000 | 0.000 | 0.000 | 0.000 | 0.000 | 0.000 | 0.000 | 0.000 | 0.000 |

**Supplementary Table 10.** Differentially abundant proteins between PD cases and controls in CSF proteomics data (limma). Proteins with adj.P.Val ≤ 0.05 and |logFC| ≥ 0.2 are shown, identified by gene symbol (HGNC) and UniProt accession, and ordered by P.Value. logFC, log2 fold change; AveExpr, average log2 abundance across samples; t, moderated t-statistic; P.Value, moderated t-test p-value; adj.P.Val, Benjamini–Hochberg adjusted p-value; B, log-odds of differential abundance.

| Protein | UniProt | logFC | AveExpr | t | P.Value | adj.P.Val | B |
|---|---|---:|---:|---:|---:|---:|---:|
| DDC | P20711 | 0.90196 | -2.03930 | 23.49745 | 5.34097E-100 | 7.81384E-97 | 214.58951 |
| PRL | P01236 | -0.98679 | 0.39575 | -11.07244 | 3.82062E-27 | 2.79478E-24 | 50.65687 |
| LPO | P22079 | -0.42666 | -2.79867 | -10.11559 | 4.23475E-23 | 2.06515E-20 | 41.57574 |
| GH1 | P01241 | 0.52153 | 0.74862 | 7.66413 | 3.81756E-14 | 1.39627E-11 | 21.53419 |
| DLK1 | P80370 | -0.22186 | -0.45547 | -7.18356 | 1.21907E-12 | 3.56700E-10 | 18.18334 |
| MMP1 | P03956 | -0.33645 | -5.45139 | -6.85129 | 1.19117E-11 | 2.90447E-9 | 15.98231 |
| GZMB | P10144 | 0.27830 | -4.69419 | 5.81767 | 7.73179E-9 | 1.61594E-6 | 9.75494 |
| TNFRSF8 | P28908 | -0.20769 | 3.00249 | -5.70180 | 1.50663E-8 | 2.75525E-6 | 9.11625 |
| CHRDL2 | Q6WN34 | -0.23778 | -3.44624 | -5.33486 | 1.15123E-7 | 1.78406E-5 | 7.17388 |
| DSG3 | P32926 | -0.20839 | -5.91373 | -5.10963 | 3.77792E-7 | 5.02463E-5 | 6.04249 |
| FCER2 | P06734 | -0.26294 | -0.64703 | -4.79434 | 1.84633E-6 | 1.71318E-4 | 4.53711 |
| CBLIF | P27352 | -0.22905 | -4.42963 | -4.79134 | 1.87361E-6 | 1.71318E-4 | 4.52322 |
| CRH | P06850 | -0.21971 | 0.43365 | -4.46705 | 8.71407E-6 | 6.05943E-4 | 3.07191 |
| VIM | P08670 | 0.32679 | 0.18095 | 4.21271 | 2.72089E-5 | 1.47432E-3 | 2.00239 |
| CEACAM21 | Q3KPI0 | -0.23231 | -2.11215 | -3.56582 | 3.77651E-4 | 1.10372E-2 | -0.44316 |
| WIF1 | Q9Y5W5 | -0.23924 | 2.63989 | -3.54177 | 4.13388E-4 | 1.11997E-2 | -0.52641 |
| CRIPTO | P13385 | -0.22440 | 0.66694 | -3.47776 | 5.24547E-4 | 1.30561E-2 | -0.74535 |
| GCG | P01275 | 0.46284 | -5.65751 | 3.44015 | 6.02276E-4 | 1.37677E-2 | -0.87216 |
| EPHA10 | Q5JZY3 | -0.20018 | 3.73570 | -3.42088 | 6.46157E-4 | 1.43231E-2 | -0.93663 |
| EPHB6 | O15197 | -0.20496 | 1.61498 | -3.28382 | 1.05500E-3 | 1.96094E-2 | -1.38480 |
| FABP1 | P07148 | -0.25848 | -3.34938 | -3.25590 | 1.16336E-3 | 2.01286E-2 | -1.47389 |
| MUC16 | Q8WXI7 | 0.36161 | -2.41243 | 2.99818 | 2.77443E-3 | 3.43982E-2 | -2.26098 |
| CLEC4G | Q6UXB4 | -0.22419 | -5.35456 | -2.97445 | 2.99653E-3 | 3.58690E-2 | -2.33025 |
| SUGT1 | Q9Y2Z0 | 0.22006 | -0.68858 | 2.97015 | 3.03847E-3 | 3.58690E-2 | -2.34275 |
| CCL13 | Q99616 | -0.28922 | -8.28735 | -2.81901 | 4.89998E-3 | 4.87665E-2 | -2.77054 |

**Supplementary Table 11.** Differentially abundant proteins between PD cases and controls in plasma proteomics data (limma). Proteins with adj.P.Val ≤ 0.05 and |logFC| ≥ 0.2 are shown, identified by gene symbol (HGNC) and UniProt accession, and ordered by P.Value. logFC, log2 fold change; AveExpr, average log2 abundance across samples; t, moderated t-statistic; P.Value, moderated t-test p-value; adj.P.Val, Benjamini–Hochberg adjusted p-value; B, log-odds of differential abundance.

| Protein | UniProt | logFC | AveExpr | t | P.Value | adj.P.Val | B |
|---|---|---|---|---|---|---|---|
| DDC | P20711 | 0.89180 | 1.36853 | 17.87422 | 2.04359E-63 | 2.98977E-60 | 132.27670 |
| PRL | P01236 | -0.88843 | -0.31967 | -11.30268 | 3.38457E-28 | 2.47582E-25 | 53.01681 |
| HPGDS | O60760 | -0.24214 | 0.34587 | -8.79324 | 5.05401E-18 | 2.46467E-15 | 30.19683 |
| SMAD1 | Q15797 | 0.62368 | 1.36272 | 8.19124 | 6.66762E-16 | 2.43868E-13 | 25.45661 |
| LDLR | P01130 | -0.28563 | -0.82744 | -8.04656 | 2.06172E-15 | 6.03259E-13 | 24.36182 |
| IL16 | Q14005 | -0.20212 | 0.51260 | -6.60916 | 5.84096E-11 | 1.22076E-8 | 14.45021 |
| ARHGAP25 | P42331 | 0.23470 | 0.57558 | 6.27115 | 5.02043E-10 | 8.16098E-8 | 12.38023 |
| CCL27 | Q9Y4X3 | -0.23607 | 0.98425 | -5.69525 | 1.55437E-8 | 1.89504E-6 | 9.08810 |
| NCLN | Q969V3 | -0.32010 | 0.25176 | -5.56361 | 3.26814E-8 | 3.22945E-6 | 8.37759 |
| GSTA3 | Q16772 | -0.29294 | 0.10339 | -5.40252 | 7.94436E-8 | 7.26412E-6 | 7.52961 |
| TNC | P24821 | 0.20201 | 0.68105 | 5.36376 | 9.80312E-8 | 7.54840E-6 | 7.32911 |
| ABHD14B | Q96IU4 | 0.36538 | 2.95771 | 5.32860 | 1.18493E-7 | 8.66773E-6 | 7.14840 |
| GSTA1 | P08263 | -0.30211 | -0.43954 | -5.03133 | 5.62786E-7 | 3.29342E-5 | 5.66601 |
| VMO1 | Q7Z5L0 | -0.21249 | 0.70452 | -4.79897 | 1.79918E-6 | 8.22564E-5 | 4.56406 |
| RBP2 | P50120 | -0.24404 | 0.67556 | -4.76745 | 2.09839E-6 | 9.30287E-5 | 4.41847 |
| FXN | Q16595 | 0.34017 | 2.99427 | 4.73268 | 2.48407E-6 | 1.00955E-4 | 4.25887 |
| PRDX1 | Q06830 | 0.22217 | 1.51688 | 4.68392 | 3.14121E-6 | 1.12088E-4 | 4.03701 |
| BGN | P21810 | -0.33042 | 0.26283 | -4.64970 | 3.69901E-6 | 1.25852E-4 | 3.88260 |
| VPS53 | Q5VIR6 | 0.29540 | 1.82114 | 4.62236 | 4.21188E-6 | 1.40045E-4 | 3.76001 |
| S100A4 | P26447 | 0.26961 | 1.96932 | 4.56561 | 5.50251E-6 | 1.75004E-4 | 3.50781 |
| PDLIM7 | Q9NR12 | 0.49765 | 4.53104 | 4.49985 | 7.47263E-6 | 2.14204E-4 | 3.21936 |
| IMPA1 | P29218 | 0.22600 | 1.51477 | 4.45820 | 9.05282E-6 | 2.40805E-4 | 3.03872 |
| CCS | O14618 | 0.22368 | 2.37849 | 4.45233 | 9.29971E-6 | 2.42955E-4 | 3.01340 |
| ABL1 | P00519 | 0.27111 | 2.12386 | 4.39923 | 1.18457E-5 | 2.98799E-4 | 2.78576 |
| SRC | P12931 | 0.48845 | 5.38347 | 4.36734 | 1.36817E-5 | 3.17236E-4 | 2.65032 |
| DPP7 | Q9UHL4 | 0.23476 | 1.34282 | 4.36627 | 1.37476E-5 | 3.17236E-4 | 2.64580 |
| NFATC1 | O95644 | 0.33241 | 3.22182 | 4.36576 | 1.37797E-5 | 3.17236E-4 | 2.64361 |
| AKT3 | Q9Y243 | 0.23664 | 1.15872 | 4.36418 | 1.38777E-5 | 3.17236E-4 | 2.63695 |
| TBC1D23 | Q9NUY8 | 0.38307 | 3.38547 | 4.35213 | 1.46502E-5 | 3.26311E-4 | 2.58606 |
| DCTN2 | Q13561 | 0.34050 | 2.85655 | 4.34867 | 1.48794E-5 | 3.26311E-4 | 2.57148 |
| STAMBP | O95630 | 0.29011 | 2.87097 | 4.32007 | 1.69104E-5 | 3.63823E-4 | 2.45132 |
| PPIB | P23284 | 0.37645 | 3.98264 | 4.29210 | 1.91508E-5 | 3.94615E-4 | 2.33455 |
| PRKAR1A | P10644 | 0.33868 | 2.48853 | 4.28344 | 1.98998E-5 | 4.04353E-4 | 2.29856 |
| STX4 | Q12846 | 0.28647 | 1.97316 | 4.26679 | 2.14197E-5 | 4.27578E-4 | 2.22952 |
| PPME1 | Q9Y570 | 0.31406 | 3.08796 | 4.26460 | 2.16273E-5 | 4.27578E-4 | 2.22048 |
| GMPR | P36959 | 0.25776 | 3.63448 | 4.23878 | 2.42283E-5 | 4.59935E-4 | 2.11401 |
| F11R | Q9Y624 | 0.21948 | 1.58258 | 4.23696 | 2.44228E-5 | 4.59935E-4 | 2.10651 |
| TYMP | P19971 | 0.20814 | 1.87428 | 4.23604 | 2.45215E-5 | 4.59935E-4 | 2.10273 |
| ERBIN | Q96RT1 | 0.35020 | 3.26376 | 4.22687 | 2.55267E-5 | 4.72729E-4 | 2.06508 |
| LYN | P07948 | 0.25288 | 2.93989 | 4.22273 | 2.59923E-5 | 4.75334E-4 | 2.04815 |
| MIF | P14174 | 0.34214 | 3.98917 | 4.21825 | 2.65069E-5 | 4.78761E-4 | 2.02978 |
| CLPP | Q16740 | 0.35934 | 3.38079 | 4.21339 | 2.70749E-5 | 4.83055E-4 | 2.00992 |
| CRADD | P78560 | 0.35364 | 4.30586 | 4.20062 | 2.86242E-5 | 5.04544E-4 | 1.95780 |
| ADH4 | P08319 | -0.22941 | -0.78566 | -4.19292 | 2.95996E-5 | 5.15527E-4 | 1.92642 |
| FABP1 | P07148 | -0.26264 | 0.79646 | -4.17132 | 3.25051E-5 | 5.42764E-4 | 1.83876 |
| CRKL | P46109 | 0.42113 | 4.98249 | 4.15751 | 3.45024E-5 | 5.42764E-4 | 1.78296 |
| DNMBP | Q6XZF7 | 0.24909 | 2.34862 | 4.15350 | 3.51032E-5 | 5.46340E-4 | 1.76681 |
| PVALB | P20472 | 0.35031 | 4.31379 | 4.14957 | 3.57024E-5 | 5.49817E-4 | 1.75098 |
| SNAP29 | O95721 | 0.37962 | 4.51537 | 4.13799 | 3.75256E-5 | 5.61065E-4 | 1.70439 |
| PLXNA4 | Q9HCM2 | 0.33291 | 3.85144 | 4.13525 | 3.79704E-5 | 5.61065E-4 | 1.69337 |
| DIABLO | Q9NR28 | 0.42468 | 4.29348 | 4.12995 | 3.88424E-5 | 5.61065E-4 | 1.67214 |
| MNDA | P41218 | -0.39352 | 0.48560 | -4.12947 | 3.89221E-5 | 5.61065E-4 | 1.67022 |
| SESTD1 | Q86VW0 | 0.26544 | 1.55801 | 4.12428 | 3.97987E-5 | 5.61065E-4 | 1.64940 |
| TDRKH | Q9Y2W6 | 0.29900 | 2.62468 | 4.12409 | 3.98297E-5 | 5.61065E-4 | 1.64867 |
| RWDD1 | Q9H446 | 0.30789 | 3.55527 | 4.12377 | 3.98843E-5 | 5.61065E-4 | 1.64739 |
| TJAP1 | Q5JTD0 | 0.32570 | 2.63594 | 4.10733 | 4.27891E-5 | 5.90738E-4 | 1.58167 |
| CASP2 | P42575 | 0.25975 | 1.54754 | 4.10726 | 4.28013E-5 | 5.90738E-4 | 1.58141 |
| ARHGEF12 | Q9NZN5 | 0.36941 | 4.10093 | 4.08331 | 4.73947E-5 | 6.43263E-4 | 1.48616 |
| SUSD1 | Q6UWL2 | 0.21968 | 2.64062 | 4.07891 | 4.82874E-5 | 6.48115E-4 | 1.46873 |
| CNPY4 | Q8N129 | 0.24898 | 2.36784 | 4.07232 | 4.96554E-5 | 6.60416E-4 | 1.44264 |
| DBNL | Q9UJU6 | 0.38687 | 5.00868 | 4.06654 | 5.08846E-5 | 6.69344E-4 | 1.41981 |

| Protein | UniProt | logFC | AveExpr | t | P.Value | adj.P.Val | B |
|---|---|---|---|---|---|---|---|
| STK11 | Q15831 | 0.27505 | 2.47863 | 4.06488 | 5.12417E-5 | 6.69344E-4 | 1.41328 |
| HTRA2 | O43464 | 0.22395 | 1.98605 | 4.06109 | 5.20705E-5 | 6.69739E-4 | 1.39830 |
| TBC1D5 | Q92609 | 0.32567 | 3.28812 | 4.02677 | 6.01606E-5 | 7.52265E-4 | 1.26352 |
| TRIM5 | Q9C035 | 0.26306 | 2.30388 | 3.97820 | 7.36714E-5 | 8.83452E-4 | 1.07464 |
| CC2D1A | Q6P1N0 | 0.33489 | 2.98722 | 3.97279 | 7.53415E-5 | 8.85517E-4 | 1.05376 |
| IRAG2 | Q12912 | 0.38055 | 2.79484 | 3.97114 | 7.58587E-5 | 8.85517E-4 | 1.04738 |
| RHOC | P08134 | 0.39441 | 5.43157 | 3.96726 | 7.70888E-5 | 8.85517E-4 | 1.03240 |
| SERPINB5 | P36952 | 0.29712 | -0.36765 | 3.96598 | 7.74966E-5 | 8.85517E-4 | 1.02748 |
| TACC3 | Q9Y6A5 | 0.41799 | 4.44883 | 3.96417 | 7.80804E-5 | 8.85517E-4 | 1.02049 |
| MAVS | Q7Z434 | 0.36560 | 3.24205 | 3.95025 | 8.27000E-5 | 9.23588E-4 | 0.96696 |
| PLPBP | O94903 | 0.28811 | 3.27334 | 3.94302 | 8.52008E-5 | 9.44309E-4 | 0.93923 |
| NCF2 | P19878 | -0.25081 | -0.12884 | -3.93811 | 8.69380E-5 | 9.56318E-4 | 0.92044 |
| ENO1 | P06733 | 0.29614 | 3.58617 | 3.93012 | 8.98414E-5 | 9.66456E-4 | 0.88986 |
| NUB1 | Q9Y5A7 | 0.29350 | 2.60274 | 3.92645 | 9.12022E-5 | 9.73933E-4 | 0.87587 |
| MESD | Q14696 | 0.41137 | 3.75560 | 3.91179 | 9.68447E-5 | 9.97773E-4 | 0.82002 |
| ANXA4 | P09525 | 0.26591 | 2.57688 | 3.90970 | 9.76793E-5 | 9.99334E-4 | 0.81204 |
| DAB2 | P98082 | 0.34383 | 3.82472 | 3.90670 | 9.88828E-5 | 1.00036E-3 | 0.80065 |
| PTPN1 | P18031 | 0.29727 | 3.02019 | 3.90604 | 9.91466E-5 | 1.00036E-3 | 0.79817 |
| BIN2 | Q9UBW5 | 0.37927 | 3.50590 | 3.90023 | 1.01525E-4 | 1.01734E-3 | 0.77613 |
| ANXA3 | P12429 | 0.26348 | 2.10064 | 3.89647 | 1.03095E-4 | 1.02604E-3 | 0.76186 |
| PPP1R12A | O14974 | 0.36250 | 3.74601 | 3.89341 | 1.04390E-4 | 1.03109E-3 | 0.75026 |
| MAP2K6 | P52564 | 0.35690 | 3.89615 | 3.87496 | 1.12512E-4 | 1.09010E-3 | 0.68061 |
| STX8 | Q9UNK0 | 0.23272 | 0.95119 | 3.87005 | 1.14775E-4 | 1.09275E-3 | 0.66210 |
| MAX | P61244 | 0.35942 | 2.64057 | 3.85620 | 1.21386E-4 | 1.13521E-3 | 0.61008 |
| SCAMP3 | O14828 | 0.32423 | 3.27320 | 3.85531 | 1.21824E-4 | 1.13521E-3 | 0.60673 |
| IPCEF1 | Q8WWN9 | 0.30458 | 3.07427 | 3.85048 | 1.24222E-4 | 1.15024E-3 | 0.58862 |
| EIF4G1 | Q04637 | 0.36666 | 4.74982 | 3.84004 | 1.29556E-4 | 1.18205E-3 | 0.54959 |
| FKBP5 | Q13451 | 0.31448 | 4.16712 | 3.83509 | 1.32160E-4 | 1.18205E-3 | 0.53112 |
| CA13 | Q8N1Q1 | 0.36958 | 5.48585 | 3.83293 | 1.33314E-4 | 1.18205E-3 | 0.52304 |
| GP6 | Q9HCN6 | 0.28957 | 2.88185 | 3.82698 | 1.36534E-4 | 1.20330E-3 | 0.50089 |
| AXIN1 | O15169 | 0.31898 | 3.80097 | 3.81581 | 1.42779E-4 | 1.24988E-3 | 0.45939 |
| HARS1 | P12081 | 0.26606 | 3.50115 | 3.81451 | 1.43527E-4 | 1.24988E-3 | 0.45454 |
| DCTN1 | Q14203 | 0.23924 | 2.81572 | 3.81265 | 1.44595E-4 | 1.25173E-3 | 0.44766 |
| VPS37A | Q8NEZ2 | 0.23897 | 2.20379 | 3.80746 | 1.47624E-4 | 1.26565E-3 | 0.42843 |
| YES1 | P07947 | 0.33913 | 3.76315 | 3.80548 | 1.48799E-4 | 1.26565E-3 | 0.42108 |
| RILP | Q96NA2 | 0.27621 | 3.88331 | 3.80085 | 1.51574E-4 | 1.27444E-3 | 0.40394 |
| COMT | P21964 | 0.28942 | 2.81986 | 3.79566 | 1.54738E-4 | 1.28828E-3 | 0.38478 |
| METAP2 | P50579 | 0.22895 | 2.48539 | 3.79527 | 1.54981E-4 | 1.28828E-3 | 0.38332 |
| METAP1D | Q6UB28 | 0.28538 | 2.58945 | 3.78931 | 1.58701E-4 | 1.30438E-3 | 0.36133 |
| CD69 | Q07108 | 0.33572 | 3.39806 | 3.78455 | 1.61728E-4 | 1.32183E-3 | 0.34382 |
| SF3B4 | Q15427 | 0.27409 | 2.44504 | 3.77836 | 1.65752E-4 | 1.33975E-3 | 0.32103 |
| CORO1A | P31146 | 0.38889 | 3.95435 | 3.77106 | 1.70615E-4 | 1.36093E-3 | 0.29424 |
| CALCOCO1 | Q9P1Z2 | 0.39235 | 5.09779 | 3.76972 | 1.71526E-4 | 1.36093E-3 | 0.28930 |
| NSFL1C | Q9UNZ2 | 0.29323 | 3.24213 | 3.76948 | 1.71690E-4 | 1.36093E-3 | 0.28842 |
| NPM1 | P06748 | -0.26550 | 1.55330 | -3.76379 | 1.75598E-4 | 1.37510E-3 | 0.26756 |
| FABP2 | P12104 | -0.21497 | 0.55673 | -3.76263 | 1.76400E-4 | 1.37510E-3 | 0.26334 |
| EBAG9 | O00559 | 0.31160 | 3.49994 | 3.76220 | 1.76704E-4 | 1.37510E-3 | 0.26174 |
| SNAP23 | O00161 | 0.32251 | 5.06313 | 3.75221 | 1.83811E-4 | 1.42283E-3 | 0.22522 |
| STK4 | Q13043 | 0.25968 | 2.63577 | 3.74172 | 1.91564E-4 | 1.45968E-3 | 0.18696 |
| CIAPIN1 | Q6FI81 | 0.26383 | 3.33006 | 3.73804 | 1.94357E-4 | 1.47261E-3 | 0.17356 |
| PRDX5 | P30044 | 0.27049 | 3.52408 | 3.73484 | 1.96809E-4 | 1.47468E-3 | 0.16195 |
| MYO9B | Q13459 | 0.27775 | 1.55700 | 3.72516 | 2.04425E-4 | 1.51814E-3 | 0.12681 |
| SKAP2 | O75563 | 0.39156 | 4.92240 | 3.71228 | 2.14995E-4 | 1.58059E-3 | 0.08016 |
| CD2AP | Q9Y5K6 | 0.22650 | 1.68033 | 3.70291 | 2.23001E-4 | 1.59147E-3 | 0.04635 |
| ATOX1 | O00244 | 0.23057 | 3.00898 | 3.69884 | 2.26559E-4 | 1.60868E-3 | 0.03171 |
| LAT2 | Q9GZY6 | 0.29869 | 2.54077 | 3.69503 | 2.29945E-4 | 1.61736E-3 | 0.01799 |
| MANF | P55145 | 0.42623 | 5.91430 | 3.69151 | 2.33116E-4 | 1.63181E-3 | 0.00533 |
| APRT | P07741 | 0.21313 | 3.89342 | 3.68297 | 2.40977E-4 | 1.67881E-3 | -0.02533 |
| NT5C3A | Q9H0P0 | 0.37050 | 4.50713 | 3.67651 | 2.47089E-4 | 1.71323E-3 | -0.04848 |
| TRIM21 | P19474 | 0.22110 | 2.86300 | 3.65622 | 2.67243E-4 | 1.82700E-3 | -0.12092 |
| DOK2 | O60496 | 0.37690 | 4.31099 | 3.64916 | 2.74603E-4 | 1.85993E-3 | -0.14601 |
| FIS1 | Q9Y3D6 | 0.24669 | 2.34681 | 3.64305 | 2.81133E-4 | 1.89538E-3 | -0.16770 |
| MAP3K5 | Q99683 | 0.39232 | 4.25266 | 3.61220 | 3.16384E-4 | 2.07565E-3 | -0.27670 |
| MGLL | Q99685 | 0.29026 | 4.36037 | 3.60530 | 3.24814E-4 | 2.11202E-3 | -0.30096 |

| Protein | UniProt | logFC | AveExpr | t | P.Value | adj.P.Val | B |
|---|---|---|---|---|---|---|---|
| SAMD9L | Q8IVG5 | 0.25063 | 2.73580 | 3.60258 | 3.28198E-4 | 2.12457E-3 | -0.31051 |
| BAX | Q07812 | 0.27823 | 2.40896 | 3.60129 | 3.29817E-4 | 2.12565E-3 | -0.31505 |
| CHMP1A | Q9HD42 | 0.25829 | 3.58727 | 3.58057 | 3.56794E-4 | 2.25969E-3 | -0.38751 |
| EIF4EBP1 | Q13541 | 0.20388 | 2.45804 | 3.56806 | 3.74071E-4 | 2.35220E-3 | -0.43107 |
| DARS1 | P14868 | 0.27754 | 3.58066 | 3.54402 | 4.09500E-4 | 2.52784E-3 | -0.51437 |
| CRACR2A | Q9BSW2 | 0.32947 | 3.12338 | 3.51934 | 4.49115E-4 | 2.70520E-3 | -0.59932 |
| CETN2 | P41208 | 0.27589 | 1.81622 | 3.51327 | 4.59394E-4 | 2.74282E-3 | -0.62012 |
| AK1 | P00568 | 0.21123 | 1.67941 | 3.51262 | 4.60494E-4 | 2.74282E-3 | -0.62232 |
| GYS1 | P13807 | 0.29863 | 3.78491 | 3.51112 | 4.63073E-4 | 2.74282E-3 | -0.62746 |
| PRDX3 | P30048 | 0.21927 | 1.69184 | 3.50032 | 4.82043E-4 | 2.80968E-3 | -0.66436 |
| SNX9 | Q9Y5X1 | 0.22417 | 2.83622 | 3.49073 | 4.99496E-4 | 2.89985E-3 | -0.69703 |
| MPI | P34949 | 0.25782 | 3.03412 | 3.48378 | 5.12519E-4 | 2.94045E-3 | -0.72067 |
| ICA1 | Q05084 | 0.24054 | 2.29753 | 3.47918 | 5.21303E-4 | 2.97917E-3 | -0.73628 |
| FADD | Q13158 | 0.26216 | 2.58101 | 3.47653 | 5.26437E-4 | 2.99437E-3 | -0.74527 |
| MEP1B | Q16820 | -0.25874 | 0.13836 | -3.47569 | 5.28056E-4 | 2.99437E-3 | -0.74810 |
| CLEC1B | Q9P126 | 0.30536 | 3.25212 | 3.47121 | 5.36872E-4 | 3.03260E-3 | -0.76330 |
| IGFBP1 | P08833 | -0.26829 | 0.93976 | -3.46261 | 5.54129E-4 | 3.09424E-3 | -0.79233 |
| STIP1 | P31948 | 0.24737 | 2.61465 | 3.46043 | 5.58594E-4 | 3.10731E-3 | -0.79970 |
| GOPC | Q9HD26 | 0.31330 | 3.74953 | 3.45153 | 5.77152E-4 | 3.19838E-3 | -0.82968 |
| SULT1A1 | P50225 | 0.37397 | 5.60834 | 3.44468 | 5.91831E-4 | 3.23078E-3 | -0.85272 |
| COX5B | P10606 | 0.20489 | 1.38892 | 3.43798 | 6.06522E-4 | 3.28645E-3 | -0.87520 |
| WASF1 | Q92558 | 0.25924 | 2.20332 | 3.42961 | 6.25350E-4 | 3.36356E-3 | -0.90323 |
| VTA1 | Q9NP79 | 0.26823 | 2.27898 | 3.42670 | 6.32025E-4 | 3.38701E-3 | -0.91296 |
| EDAR | Q9UNE0 | 0.23648 | 1.92259 | 3.42446 | 6.37205E-4 | 3.40230E-3 | -0.92044 |
| GH1 | P01241 | 0.40786 | 3.29138 | 3.41110 | 6.68917E-4 | 3.55864E-3 | -0.96494 |
| SPRY2 | O43597 | 0.26572 | 2.56063 | 3.40768 | 6.77275E-4 | 3.57709E-3 | -0.97632 |
| BCR | P11274 | 0.25780 | 3.26952 | 3.38957 | 7.23170E-4 | 3.79210E-3 | -1.03635 |
| PSG1 | P11464 | 0.25615 | -0.97748 | 3.38792 | 7.27507E-4 | 3.80123E-3 | -1.04182 |
| STK24 | Q9Y6E0 | 0.22407 | 1.44682 | 3.38186 | 7.43563E-4 | 3.83126E-3 | -1.06180 |
| KIFBP | Q96EK5 | 0.26910 | 3.00319 | 3.38180 | 7.43731E-4 | 3.83126E-3 | -1.06200 |
| FYB1 | O15117 | 0.30487 | 4.86726 | 3.37297 | 7.67752E-4 | 3.94113E-3 | -1.09109 |
| ITGB1BP2 | Q9UKP3 | 0.33554 | 4.60131 | 3.36591 | 7.87497E-4 | 4.00533E-3 | -1.11431 |
| DNAJA2 | O60884 | 0.27617 | 4.14760 | 3.36550 | 7.88654E-4 | 4.00533E-3 | -1.11565 |
| IKZF2 | Q9UKS7 | -0.29629 | 0.45296 | -3.36460 | 7.91210E-4 | 4.00533E-3 | -1.11861 |
| ENO2 | P09104 | 0.21955 | 2.80974 | 3.35686 | 8.13463E-4 | 4.10378E-3 | -1.14397 |
| NCK2 | O43639 | 0.27042 | 2.87567 | 3.35326 | 8.24009E-4 | 4.11622E-3 | -1.15575 |
| IKBKG | Q9Y6K9 | 0.25738 | 3.46140 | 3.35314 | 8.24369E-4 | 4.11622E-3 | -1.15615 |
| HPCAL1 | P37235 | 0.22582 | 3.50076 | 3.34315 | 8.54348E-4 | 4.18357E-3 | -1.18879 |
| CNST | Q6PJW8 | 0.28526 | 3.63257 | 3.34293 | 8.55015E-4 | 4.18357E-3 | -1.18950 |
| LACTB2 | Q53H82 | 0.21687 | 2.55939 | 3.33949 | 8.65572E-4 | 4.22111E-3 | -1.20071 |
| DBI | P07108 | 0.22307 | 2.64532 | 3.33559 | 8.77689E-4 | 4.25185E-3 | -1.21341 |
| CLIP2 | Q9UDT6 | 0.33676 | 5.05649 | 3.32925 | 8.97709E-4 | 4.27833E-3 | -1.23401 |
| LAT | O43561 | 0.24826 | 4.13596 | 3.32034 | 9.26610E-4 | 4.40140E-3 | -1.26294 |
| CASP3 | P42574 | 0.29077 | 5.02012 | 3.31641 | 9.39618E-4 | 4.43439E-3 | -1.27567 |
| GRAP2 | O75791 | 0.37206 | 5.02236 | 3.30752 | 9.69666E-4 | 4.56148E-3 | -1.30439 |
| CDC37 | Q16543 | 0.26264 | 2.29662 | 3.28427 | 1.05249E-3 | 4.85741E-3 | -1.37914 |
| PPP1R2 | P41236 | 0.21799 | 2.58449 | 3.25451 | 1.16814E-3 | 5.32396E-3 | -1.47412 |
| MPIG6B | O95866 | 0.24538 | 3.43852 | 3.24694 | 1.19937E-3 | 5.39899E-3 | -1.49813 |
| GZMH | P20718 | -0.21813 | -0.31197 | -3.21479 | 1.34083E-3 | 5.90855E-3 | -1.59955 |
| CDKN2D | P55273 | 0.29864 | 3.96788 | 3.21111 | 1.35796E-3 | 5.94821E-3 | -1.61109 |
| TXLNA | P40222 | 0.23731 | 3.74390 | 3.20755 | 1.37472E-3 | 5.98575E-3 | -1.62223 |
| FHIT | P49789 | 0.27543 | 3.19740 | 3.20587 | 1.38272E-3 | 6.00272E-3 | -1.62750 |
| HEXIM1 | O94992 | 0.23351 | 3.60585 | 3.19515 | 1.43466E-3 | 6.18204E-3 | -1.66100 |
| CASP10 | Q92851 | 0.22645 | 1.83704 | 3.19473 | 1.43670E-3 | 6.18204E-3 | -1.66229 |
| CD40LG | P29965 | 0.23719 | 1.96712 | 3.18935 | 1.46350E-3 | 6.24229E-3 | -1.67908 |
| MGMT | P16455 | 0.32751 | 4.83724 | 3.13852 | 1.74014E-3 | 7.23244E-3 | -1.83610 |
| EGF | P01133 | 0.29007 | 3.24608 | 3.13602 | 1.75497E-3 | 7.27343E-3 | -1.84379 |
| DECR1 | Q16698 | 0.25866 | 3.93803 | 3.12576 | 1.81683E-3 | 7.46637E-3 | -1.87517 |
| PLA2G4A | P47712 | 0.24019 | 3.05432 | 3.11629 | 1.87571E-3 | 7.61954E-3 | -1.90404 |
| MSRA | Q9UJ68 | 0.20340 | 2.73856 | 3.11570 | 1.87940E-3 | 7.61954E-3 | -1.90582 |
| SERPINB1 | P30740 | 0.22588 | 2.66315 | 3.11314 | 1.89564E-3 | 7.64000E-3 | -1.91360 |
| STAT5B | P51692 | 0.39060 | 6.06740 | 3.10968 | 1.91783E-3 | 7.66823E-3 | -1.92414 |
| PRTFDC1 | Q9NRG1 | 0.32875 | 4.25876 | 3.10959 | 1.91837E-3 | 7.66823E-3 | -1.92439 |
| RNASE3 | P12724 | -0.28311 | 1.01772 | -3.10678 | 1.93658E-3 | 7.69897E-3 | -1.93294 |

| Protein | UniProt | logFC | AveExpr | t | P.Value | adj.P.Val | B |
|---|---|---|---|---|---|---|---|
| PFKFB2 | O60825 | 0.20852 | 3.00946 | 3.09761 | 1.99698E-3 | 7.87743E-3 | -1.96071 |
| RUVBL1 | Q9Y265 | -0.22964 | -0.11127 | -3.09751 | 1.99763E-3 | 7.87743E-3 | -1.96101 |
| MAPT | P10636 | 0.29181 | -0.37730 | 3.08621 | 2.07450E-3 | 8.11494E-3 | -1.99514 |
| TARBP2 | Q15633 | 0.23901 | 2.99110 | 3.08117 | 2.10970E-3 | 8.23062E-3 | -2.01034 |
| SIRT2 | Q8IXJ6 | 0.28000 | 4.15837 | 3.08020 | 2.11647E-3 | 8.23510E-3 | -2.01324 |
| PPP1R9B | Q96SB3 | 0.22540 | 3.45297 | 3.07523 | 2.15182E-3 | 8.35043E-3 | -2.02821 |
| DAPP1 | Q9UN19 | 0.33494 | 4.76917 | 3.06108 | 2.25526E-3 | 8.63727E-3 | -2.07060 |
| TBCB | Q99426 | 0.30398 | 5.56388 | 3.05554 | 2.29706E-3 | 8.74040E-3 | -2.08718 |
| IRAK4 | Q9NWZ3 | 0.30769 | 4.66222 | 3.05514 | 2.30011E-3 | 8.74040E-3 | -2.08838 |
| SUGT1 | Q9Y2Z0 | 0.24017 | 3.14370 | 3.05308 | 2.31580E-3 | 8.77724E-3 | -2.09451 |
| TRAF2 | Q12933 | 0.21215 | 2.87907 | 3.04467 | 2.38105E-3 | 8.96585E-3 | -2.11958 |
| MAP4K5 | Q9Y4K4 | 0.28821 | 3.64018 | 3.03778 | 2.43565E-3 | 9.09020E-3 | -2.14003 |
| INPPL1 | O15357 | 0.24085 | 3.12869 | 2.99895 | 2.76592E-3 | 1.01163E-2 | -2.25458 |
| TBCC | Q15814 | 0.22781 | 3.48108 | 2.97972 | 2.94417E-3 | 1.06880E-2 | -2.31076 |
| TCL1A | P56279 | -0.23293 | 1.73764 | -2.97192 | 3.01947E-3 | 1.08680E-2 | -2.33346 |
| BANK1 | Q8NDB2 | 0.27238 | 4.94529 | 2.95490 | 3.18979E-3 | 1.13821E-2 | -2.38275 |
| ZBTB16 | Q05516 | 0.20737 | 1.94456 | 2.94270 | 3.31728E-3 | 1.16663E-2 | -2.41793 |
| ATP5IF1 | Q9UII2 | 0.26955 | 4.75322 | 2.93738 | 3.37435E-3 | 1.18337E-2 | -2.43324 |
| EPCAM | P16422 | 0.20004 | 0.83733 | 2.93580 | 3.39148E-3 | 1.18349E-2 | -2.43778 |
| ANXA11 | P50995 | 0.20776 | 3.55361 | 2.92305 | 3.53242E-3 | 1.22237E-2 | -2.47429 |
| MITD1 | Q8WV92 | 0.25603 | 4.23030 | 2.91643 | 3.60768E-3 | 1.23870E-2 | -2.49319 |
| USO1 | O60763 | 0.24918 | 3.56236 | 2.88553 | 3.97877E-3 | 1.32596E-2 | -2.58085 |
| PTPN6 | P29350 | 0.23598 | 4.44104 | 2.82141 | 4.86158E-3 | 1.58055E-2 | -2.75981 |
| FGF21 | Q9NSA1 | -0.22922 | 1.66185 | -2.80593 | 5.09972E-3 | 1.64700E-2 | -2.80241 |
| AIFM1 | O95831 | 0.25900 | 2.69533 | 2.79241 | 5.31630E-3 | 1.70565E-2 | -2.83944 |
| EIF4B | P23588 | 0.21355 | 3.91956 | 2.77669 | 5.57845E-3 | 1.75889E-2 | -2.88225 |
| DNAJB1 | P25685 | 0.25398 | 3.57417 | 2.76348 | 5.80791E-3 | 1.82731E-2 | -2.91807 |
| SH2B3 | Q9UQQ2 | 0.23491 | 3.30685 | 2.73382 | 6.35405E-3 | 1.97367E-2 | -2.99784 |
| PDGFB | P01127 | 0.21697 | 2.87787 | 2.72569 | 6.51184E-3 | 2.00988E-2 | -3.01958 |
| FOLR3 | P41439 | -0.28214 | -2.94738 | -2.71252 | 6.77459E-3 | 2.07783E-2 | -3.05462 |
| CXCL3 | P19876 | 0.20449 | 4.62923 | 2.63149 | 8.61197E-3 | 2.51986E-2 | -3.26656 |
| CCL19 | Q99731 | 0.21245 | 1.15844 | 2.62443 | 8.79167E-3 | 2.55710E-2 | -3.28474 |
| TIA1 | P31483 | 0.25375 | 2.47406 | 2.61576 | 9.01657E-3 | 2.60787E-2 | -3.30697 |
| IL4 | P05112 | 0.25390 | -0.44095 | 2.54479 | 1.10611E-2 | 3.08591E-2 | -3.48633 |
| PMVK | Q15126 | 0.27542 | 6.92561 | 2.53375 | 1.14137E-2 | 3.15657E-2 | -3.51379 |
| USP8 | P40818 | 0.21166 | 0.73137 | 2.51012 | 1.22021E-2 | 3.34929E-2 | -3.57216 |
| FKBP1B | P68106 | 0.20537 | 1.79122 | 2.50573 | 1.23537E-2 | 3.37822E-2 | -3.58293 |
| CDKN1A | P38936 | 0.22666 | 3.76626 | 2.50449 | 1.23968E-2 | 3.38368E-2 | -3.58597 |
| GPA33 | Q99795 | 0.22396 | 1.44046 | 2.39994 | 1.65523E-2 | 4.27089E-2 | -3.83721 |

**Supplementary Table 12.** MariNET adjacency matrix for whole-blood transcriptomics. Values are the edge weights (t-values) between each of the 10 differentially expressed genes included in the network and each clinical variable. UPDRS1–4, MDS-UPDRS Parts I–IV; MOCA, Montreal Cognitive Assessment; PDQ-39 domains; Schwad_ADL, Schwab and England Activities of Daily Living; ESS, Epworth Sleepiness Scale; UPSIT, University of Pennsylvania Smell Identification Test.

| **Clinical variable** | ***TENT5C*** | ***ANKRD36 BP2*** | ***FOLR3*** | ***XK*** | ***SLC6A8*** | ***CA1*** | ***H1-0*** | ***SPTB*** | ***TRIM58*** | ***PITHD1*** |
|---|---|---|---|---|---|---|---|---|---|---|
| UPDRS1 | 0.680 | -0.648 | -2.137 | -0.928 | 0.239 | 0.711 | -0.044 | -0.348 | -0.078 | 0.227 |
| UPDRS2 | 0.102 | 0.425 | 1.294 | -0.449 | -1.007 | 0.319 | 1.031 | 0.030 | -1.099 | -0.718 |
| UPDRS3 | -1.021 | -1.614 | -1.494 | -0.409 | -0.573 | -0.330 | -1.359 | -1.351 | -0.741 | -0.950 |
| UPDRS4 | -1.451 | -1.732 | 0.471 | 0.735 | -0.081 | -1.389 | -1.558 | -0.082 | -1.227 | -1.667 |
| MOCA | 0.981 | -0.100 | 1.148 | 0.926 | 0.442 | 0.105 | 0.402 | 1.756 | 1.262 | 1.178 |
| Mobility39 | 0.578 | -0.611 | 2.012 | 0.973 | -0.482 | -0.440 | -0.636 | -0.865 | -1.289 | 0.275 |
| ADL39 | -0.883 | 0.965 | 0.575 | 0.146 | 1.212 | 0.716 | 1.173 | -0.527 | 1.683 | 0.466 |
| Emotional39 | -0.864 | -0.390 | 0.362 | -0.510 | -1.520 | -0.165 | -0.707 | 1.020 | -0.445 | -0.176 |
| Stigma39 | -0.125 | 0.030 | -0.891 | -0.986 | -0.016 | -0.897 | 1.047 | -0.497 | 0.706 | -0.469 |
| Social39 | 1.167 | 2.456 | 0.413 | 1.388 | 1.905 | 0.335 | -1.638 | -0.038 | 0.788 | 0.413 |
| Cognition39 | -0.872 | -0.226 | -1.166 | 1.135 | -0.688 | -0.575 | 0.385 | -0.289 | 0.580 | -0.941 |
| Communication39 | 1.383 | -1.097 | 0.639 | -1.260 | 0.218 | 0.465 | 1.125 | -0.141 | -0.797 | 1.105 |
| Discomfort39 | 1.081 | 0.918 | 0.111 | 2.004 | 0.437 | 0.964 | 0.033 | -0.570 | 0.419 | 1.060 |
| Schwad_ADL | 0.205 | 0.609 | 0.785 | 0.519 | 0.306 | -0.455 | 1.836 | -1.463 | -0.804 | 0.824 |
| ESS | -0.899 | -0.466 | 0.443 | 0.374 | 0.073 | -0.301 | -0.146 | 0.073 | 0.603 | 0.343 |
| UPSIT | -2.473 | 0.421 | -0.970 | -0.761 | -1.441 | -1.900 | 0.401 | -0.275 | -0.905 | -0.217 |

**Supplementary Table 13.** MariNET adjacency matrix for whole-blood transcriptomic pathways. Values are the edge weights (t-values) between each pathway included in the network (the 13 non-redundant Gene Ontology Biological Process terms in Supplementary Table 5) and each clinical variable. Clinical variable abbreviations as in Supplementary Table 12.

| Pathway | UPDRS1 | UPDRS2 | UPDRS3 | UPDRS4 | MOCA | Mobility39 | ADL39 | Emotional39 | Stigma39 | Social39 | Cognition39 | Communication39 | Discomfort39 | Schwad_ADL | ESS | UPSIT |
|---|---|---|---|---|---|---|---|---|---|---|---|---|---|---|---|---|
| Oxidative phosphorylation | -0.110 | 0.294 | -1.629 | -1.510 | 0.073 | 0.965 | 1.201 | 0.693 | -0.797 | 1.315 | -1.251 | -0.021 | 0.244 | 1.015 | -0.099 | -0.209 |
| Signal transduction | -1.073 | 0.766 | -0.542 | 0.255 | -0.807 | -0.168 | -1.423 | 0.254 | 0.186 | 0.617 | 0.309 | 0.676 | 1.039 | -0.591 | -0.795 | -1.343 |
| G protein-coupled receptor signaling pathway | 0.212 | -0.682 | -0.760 | 0.181 | 0.087 | 0.638 | 0.912 | -0.475 | -1.024 | 0.443 | -0.398 | 0.666 | -1.325 | 0.964 | -0.513 | -1.747 |
| Negative regulation of megakaryocyte differentiation | 0.633 | -1.000 | -1.068 | 0.399 | 0.208 | 0.541 | 0.624 | 0.799 | 0.378 | -0.909 | -1.418 | 1.235 | 0.701 | 0.708 | 0.637 | -1.222 |
| Intracellular signal transduction | -0.491 | -0.038 | 1.202 | 0.377 | 0.448 | 0.475 | -0.025 | 0.442 | -1.358 | 1.207 | 0.207 | -0.154 | -1.409 | -0.294 | 0.329 | -1.099 |
| Protein polyubiquitination | 0.612 | 0.248 | 0.222 | -0.752 | -1.697 | -0.552 | -0.060 | -0.365 | -1.428 | 1.528 | -0.406 | -0.390 | 1.967 | 0.620 | 1.443 | 0.211 |
| Phospholipase C-activating G protein-coupled receptor signaling pathway | 0.594 | 1.034 | -1.956 | -1.390 | -0.730 | 0.784 | -0.539 | -0.209 | 0.853 | -1.035 | -0.890 | 0.788 | -0.250 | 0.970 | -0.552 | -2.737 |
| DNA replication-dependent chromatin assembly | -0.167 | 0.712 | -0.495 | -1.547 | -0.589 | 0.522 | 0.518 | -0.540 | -1.185 | 1.371 | 0.248 | -0.664 | 0.328 | 0.428 | -0.705 | 0.440 |
| Positive regulation of T-helper 1 cell cytokine production | -0.637 | -0.464 | -1.755 | 0.292 | -2.874 | 0.774 | 1.285 | 0.382 | -1.104 | 0.884 | 0.169 | 0.462 | 0.271 | -0.071 | -1.153 | -0.626 |
| Proteasome-mediated ubiquitin-dependent protein catabolic process | -1.851 | -0.634 | -1.428 | -0.779 | -0.054 | 0.084 | 1.347 | 0.682 | -0.203 | -0.227 | -0.041 | 1.075 | 1.118 | 0.256 | -0.294 | 0.258 |
| Protein ubiquitination | -0.216 | -0.034 | -0.126 | -0.892 | -0.491 | -0.105 | -0.614 | -0.430 | -0.228 | 0.330 | 0.721 | -0.719 | 0.428 | -0.117 | -0.709 | -0.304 |
| Chemotaxis | -1.725 | -0.348 | 0.169 | 0.373 | -0.776 | -1.042 | 2.069 | 0.456 | -0.883 | 1.525 | -0.647 | 0.250 | 1.005 | 1.770 | 0.408 | -0.756 |
| Chromatin organization | -0.644 | 2.385 | 0.439 | -0.044 | 0.296 | -2.233 | -2.654 | 0.500 | 1.088 | -1.170 | 2.226 | -0.818 | -0.725 | -2.322 | -0.914 | 0.628 |

**Supplementary Table 14.** MariNET adjacency matrix for CSF proteomics. Values are the edge weights (t-values) between each of the 10 CSF differentially abundant proteins included in the network and each clinical variable. Clinical variable abbreviations as in Supplementary Table 12.

| Clinical variable | PRL | DDC | GCG | TNFRSF8 | CHRDL2 | CBLIF | EPHB6 | FABP1 | CLEC4G | EPHA10 |
|---|---|---|---|---|---|---|---|---|---|---|
| UPDRS1 | 1.352 | 0.411 | -0.275 | 1.098 | 1.743 | 2.517 | 0.139 | -0.260 | -2.001 | 0.949 |
| UPDRS2 | -0.505 | 1.310 | 0.595 | 0.370 | -0.911 | -1.673 | 0.623 | 0.701 | 0.328 | -0.778 |
| UPDRS3 | 1.041 | 0.213 | 0.723 | 0.708 | 0.928 | -0.497 | 0.571 | -0.639 | 0.137 | -0.970 |
| UPDRS4 | -1.173 | 2.793 | 1.133 | -0.298 | -0.619 | 0.016 | 0.971 | -0.329 | -0.479 | 1.077 |
| MOCA | -0.413 | 1.300 | -0.655 | -0.463 | 0.404 | -0.771 | 0.748 | -1.354 | 0.202 | 1.275 |
| Mobility39 | 0.671 | -1.141 | -2.784 | 1.912 | 1.527 | -0.367 | 1.061 | 1.739 | 0.163 | 0.410 |
| ADL39 | 0.882 | -1.741 | 0.517 | 0.322 | 0.442 | 1.493 | 0.382 | -0.982 | -0.172 | 0.700 |
| Emotional39 | 1.007 | -0.746 | -0.364 | -0.150 | -2.710 | 0.733 | -0.376 | -1.307 | 1.163 | -2.225 |
| Stigma39 | 0.105 | 1.440 | 0.985 | -2.768 | 0.205 | -0.244 | -2.307 | -0.863 | -0.864 | -0.834 |
| Social39 | -0.142 | -0.699 | 0.715 | -1.446 | -0.047 | -0.193 | -1.000 | 0.209 | -0.079 | -0.008 |
| Cognition39 | 0.623 | -0.617 | -0.839 | -0.532 | 0.798 | -2.670 | 0.381 | -0.249 | 0.655 | 0.190 |
| Communication39 | 0.578 | 0.188 | 0.286 | 0.821 | -0.645 | -1.653 | 0.101 | 0.583 | 2.225 | -0.533 |
| Discomfort39 | -0.558 | 0.686 | 1.082 | 0.028 | 0.161 | 0.226 | 0.014 | 2.237 | -1.272 | 2.109 |
| Schwad_ADL | 0.673 | -2.521 | 0.405 | -0.272 | 1.113 | -0.788 | 1.153 | 0.291 | 0.000 | -0.134 |
| ESS | -2.847 | 0.966 | 0.736 | 0.382 | -0.141 | 0.294 | -0.172 | 1.373 | -0.189 | 1.208 |
| UPSIT | -0.452 | -0.558 | -0.038 | -1.859 | 1.551 | 0.210 | -0.258 | -1.360 | 0.198 | -0.050 |

**Supplementary Table 15.** MariNET adjacency matrix for plasma proteomics. Values are the edge weights (t-values) between each of the 10 plasma differentially abundant proteins included in the network and each clinical variable. Clinical variable abbreviations as in Supplementary Table 12.

| **Clinical variable** | **SF3B4** | **DDC** | **RNASE3** | **CLEC1B** | **FGF21** | **CDKN2D** | **STK11** | **PRDX1** | **ADH4** | **CC2D1A** |
|---|---|---|---|---|---|---|---|---|---|---|
| UPDRS1 | -0.348 | -0.137 | 0.183 | -0.403 | -0.050 | 0.683 | 0.954 | -0.908 | 0.146 | -0.355 |
| UPDRS2 | -1.073 | 0.921 | -0.123 | -0.165 | -0.911 | -0.896 | -0.097 | -0.945 | -0.029 | -1.066 |
| UPDRS3 | 0.698 | -0.437 | -1.130 | 0.149 | -1.362 | -0.197 | 2.415 | 1.156 | -0.677 | 0.284 |
| UPDRS4 | -0.229 | 2.798 | -1.579 | 0.771 | 0.035 | -0.980 | 0.133 | -0.312 | -0.915 | 0.026 |
| MOCA | 0.985 | -0.291 | -0.288 | -0.269 | 0.440 | 0.518 | 0.185 | 0.222 | 0.218 | 0.137 |
| Mobility39 | 1.342 | -0.850 | 1.483 | 0.925 | 0.670 | 2.437 | 0.041 | 1.828 | 0.147 | 0.993 |
| ADL39 | 0.655 | -1.192 | -0.310 | 0.280 | 0.782 | 0.140 | -0.567 | 0.519 | 1.815 | 0.656 |
| Emotional39 | -1.372 | -0.361 | 2.606 | -0.325 | -0.708 | -0.423 | 0.201 | -0.168 | -2.321 | 0.886 |
| Stigma39 | 0.897 | 1.336 | -0.589 | 0.194 | 0.065 | 0.147 | 0.000 | -0.863 | 0.972 | -0.915 |
| Social39 | -0.218 | 0.278 | -0.089 | 0.105 | 0.792 | 0.217 | -0.396 | 0.200 | 0.655 | -0.059 |
| Cognition39 | 1.176 | -0.560 | -0.318 | 2.550 | 0.238 | 0.712 | 0.392 | 2.349 | 0.259 | 2.283 |
| Communication39 | -1.684 | -0.541 | 0.486 | -2.009 | 1.196 | 0.853 | -0.460 | -0.721 | -1.581 | -0.557 |
| Discomfort39 | -0.475 | 0.734 | -1.870 | -0.339 | -1.090 | -2.371 | -0.977 | -0.924 | 0.767 | -0.687 |
| Schwad_ADL | 0.167 | -0.443 | 0.982 | -0.072 | 0.334 | 1.034 | 0.706 | -0.666 | 0.602 | -0.636 |
| ESS | -0.547 | 0.025 | -0.876 | -0.314 | -2.461 | -0.716 | 1.199 | -0.018 | -1.422 | -0.442 |
| UPSIT | 2.964 | -0.387 | -0.559 | 1.283 | -0.369 | 0.002 | 0.191 | 1.615 | 1.139 | 1.349 |

**Supplementary Table 16.** Comparison between CSF and plasma of the MariNET edge weights (t-values) of the five differentially abundant proteins shared between both compartments. For each protein, values from the CSF network and the plasma network are shown side by side for every clinical variable. Clinical variable abbreviations as in Supplementary Table 12.

| | DDC | | PRL | | FABP1 | | GH1 | | SUGT1 | |
|---|---|---|---|---|---|---|---|---|---|---|
| **Clinical variable** | **CSF** | **Plasma** | **CSF** | **Plasma** | **CSF** | **Plasma** | **CSF** | **Plasma** | **CSF** | **Plasma** |
| UPDRS1 | 0.761 | 0.196 | 1.633 | 1.013 | 0.638 | -0.371 | 0.779 | -0.189 | -0.515 | 0.486 |
| UPDRS2 | 1.092 | 0.891 | -0.546 | -0.462 | 0.341 | -0.426 | -0.253 | -0.944 | 0.045 | 0.630 |
| UPDRS3 | 0.186 | -1.099 | 1.097 | 1.081 | -0.705 | -0.882 | 1.439 | 1.629 | -0.106 | 2.247 |
| UPDRS4 | 2.661 | 3.560 | -1.299 | -1.204 | -1.123 | -1.244 | 0.791 | -0.232 | -0.388 | -0.042 |
| MOCA | 1.441 | 0.503 | -0.329 | -0.515 | -1.619 | -0.694 | 1.618 | 0.269 | -1.209 | -0.783 |
| Mobility39 | -0.773 | -0.474 | 1.032 | 1.123 | 2.472 | 2.124 | 1.391 | -1.012 | 0.797 | 1.146 |
| ADL39 | -1.506 | -2.027 | 1.127 | 0.866 | -0.483 | -0.546 | -0.314 | 1.344 | -0.938 | -1.011 |
| Emotional39 | -1.100 | -0.296 | 1.249 | 2.221 | -1.435 | -0.580 | -1.312 | -0.418 | 1.025 | 0.125 |
| Stigma39 | 1.194 | 2.006 | -0.545 | -0.337 | -2.032 | -1.052 | 0.726 | 1.564 | 0.628 | 0.180 |
| Social39 | -0.798 | -0.081 | -0.529 | -0.713 | 0.171 | 0.750 | -0.058 | 1.097 | -1.416 | 0.148 |
| Cognition39 | -0.470 | 0.364 | 0.331 | 0.294 | -0.942 | -1.094 | -1.166 | 0.912 | 1.052 | 0.664 |
| Communication39 | -0.006 | -0.563 | 0.579 | -0.612 | 0.158 | -0.093 | 0.625 | -0.461 | 0.099 | -0.235 |
| Discomfort39 | 0.411 | 0.687 | -0.732 | -1.006 | 2.452 | 1.071 | -0.046 | -1.025 | 0.682 | -1.321 |
| Schwad_ADL | -2.367 | -1.021 | 0.683 | 0.377 | 0.775 | 1.458 | -0.452 | -0.354 | 0.456 | 0.740 |
| ESS | 0.746 | 0.680 | -3.158 | -3.078 | 1.372 | 1.644 | 0.871 | 0.775 | -0.952 | -0.206 |
| UPSIT | 0.048 | -0.296 | -0.632 | -0.322 | -1.360 | -2.092 | -1.548 | 0.445 | 0.264 | -1.181 |

**Supplementary Table 17.** Edges between the five differentially abundant proteins shared between CSF and plasma and the clinical variables with an absolute edge weight (|t|) greater than 1 in both the CSF and the plasma networks. Pearson correlation between CSF and plasma edge weights: r = 0.88 for the edges shown and r = 0.68 for all protein–clinical edges (Supplementary Table 16). Clinical variable abbreviations as in Supplementary Table 12.

| Protein | Clinical variable | t-value (CSF) | t-value (plasma) |
|---|---|---:|---:|
| DDC | UPDRS4 | 2.661 | 3.560 |
| | ADL39 | -1.506 | -2.027 |
| | Stigma39 | 1.194 | 2.006 |
| | Schwad_ADL | -2.367 | -1.021 |
| PRL | UPDRS1 | 1.633 | 1.013 |
| | UPDRS3 | 1.097 | 1.081 |
| | UPDRS4 | -1.299 | -1.204 |
| | Mobility39 | 1.032 | 1.123 |
| | Emotional39 | 1.249 | 2.221 |
| | ESS | -3.158 | -3.078 |
| FABP1 | UPDRS4 | -1.123 | -1.244 |
| | Mobility39 | 2.472 | 2.124 |
| | Stigma39 | -2.032 | -1.052 |
| | Discomfort39 | 2.452 | 1.071 |
| | ESS | 1.372 | 1.644 |
| | UPSIT | -1.360 | -2.092 |
| GH1 | UPDRS3 | 1.439 | 1.629 |
| | Mobility39 | 1.391 | -1.012 |
